# Convergence and molecular evolution of floral fragrance after independent transitions to self–fertilization

**DOI:** 10.1101/2022.10.04.510758

**Authors:** Natalia Wozniak, Kevin Sartori, Christian Kappel, Lihua Zhao, Alexander Erban, Ines Fehrle, Friederike Jantzen, Marion Orsucci, Stefanie Rosa, Michael Lenhard, Joachim Kopka, Adrien Sicard

## Abstract

The study of the independent evolution of similar characters can highlight important ecological and genetic factors that drive phenotypic evolution. The transition from reproduction by outcrossing to self-fertilization has occurred frequently throughout plant evolution. A common trend in this transition is the reduction of flower features in the selfing lineages, including display size, flower signals and pollinators’ rewards. These changes are believed to evolve because resources invested in building attractive flowers are reallocated to other fitness functions as the pressures to attract pollinators decrease. We investigated the similarities in the evolution of flower fragrance after independent transitions to self-fertilization in *Capsella*. We identified a large number of compounds that are similarly changed in different selfer lineages, such that the composition of the flower scent can predict the mating system in this genus. We further demonstrate that the emission of some of these compounds convergently evolved based on mutations in different genes. In one of the *Capsella* selfing lineages, the loss of *β*-ocimene emission was caused by a mutation altering subcellular localization of the ortholog of TERPENE SYNTHASE 2 without apparent effects on its biosynthetic activity. This mutation appears to have been selected at the early stage of this selfing lineage establishment through the capture of a variant segregating in the ancestral outcrossing population. The large extent of convergence in the independent evolution of flower scent, together with the evolutionary history and molecular consequences of a causal mutation, suggest that the emission of specific volatiles has important fitness consequences in self-fertilizing plants without obvious energetic benefits.

## Introduction

Understanding the factors that influence evolutionary trajectories is essential to predicting future changes in biodiversity. The convergent evolution of similar phenotypes is particularly informative because it represents a natural replay-experiment that can reveal the ecological and genetic contexts leading to evolutionary repeatability (Blount *et al*., 2018). In plants, one of the most common repeated evolutionary events is the shift from outbreeding to self-fertilization. With 10 to 15% of flowering plants being predominantly selfing, this transition is pervasive during plant evolution and has occurred at least once in most angiosperm genera (Stebbins, 1957). It is generally initiated by a mutation inactivating self-incompatibility systems in outcrossing lineages, which prevent pollen grains from fertilizing ovules from the same plant genotype (Takayama and Isogai, 2005; Wright *et al*., 2013; Durand *et al*., 2020). To establish a new selfing lineage, such mutations must provide a reproductive assurance advantage that outweighs the cost of inbreeding depression (Fisher, 1941; Busch and Delph, 2012). Strikingly, selfing lineages display very similar changes in flower morphology and function, including a reduction in petal size, a change in the resource allocation to sexual function and a decrease in the production of nectar and scent (Sicard and Lenhard, 2011). The similarities in the trajectory of flower evolution between independently evolved selfers are such that this suite of change has been termed the ‘selfing syndrome’. This also implies that the mode of reproduction highly influences the intensity and direction of the selective pressure imposed on floral structure and function.

Because the reproductive success of obligate outcrossing species is entirely dependent on pollinators, intense selection promotes the maintenance of attractive floral features (Yan *et al*., 2016). As a result, outcrossing species often develop large and showy flowers offering large rewards to pollinators (Sicard and Lenhard, 2011). In addition, they also emit a complex mixture of volatile chemical compounds believed to influence the rate of pollinator visitation (Knudsen and Gershenzon, 2006; Schiestl and Johnson, 2013). However, because in selfing lineages pollinators are not required to reproduce, the pressures to maintain these attractive features are likely released, leading to the evolution of the selfing syndrome (Sicard and Lenhard, 2011). Decreases in flower size, pollen number, nectar production, and volatile emission are, therefore, often assumed to evolve as the result of the relaxation of selective pressure or because resource limitations shift energy allocation to other fitness related function, thereby selecting against costly flower phenotypes (Barrett, 2002; Sicard and Lenhard, 2011). Considering the stochastic nature of mutations, these theories imply that different selfing syndrome traits should evolve differently among selfing lineages depending on the time of selfing evolution. Conversely, some of these traits may provide fitness advantages in selfers, by acting as mating system modifiers, improving self-fertilization or avoiding reproductive interference through pollinator avoidance (Barrett, 2013; Orsucci and Sicard, 2021). The selection of such traits could therefore promote a high level of similarities in independent transitions to self-fertilization. In addition to this fitness argument, the structure of the gene regulatory networks controlling these traits may also limit the number of fitness-neutral or advantageous mutations in the genomes that can lead to the evolution of the selfing syndrome, thereby increasing the level of phenotypic convergence (Wozniak *et al*., 2020). Therefore, comparing the independent evolution of self-fertilization can be used to highlight traits with high reproductive fitness for different mating systems and/or the existence of constraints driving phenotypic evolution. Studies that compared independent occurrences of the selfing syndrome found high levels of phenotypic and molecular convergence in flower morphology (Wozniak *et al*., 2020). However, the relationship between mating system shifts and the evolution of scent emissions has not been extensively studied. So far, it has been shown that selfing populations are associated with shifts in the composition of floral scent but not necessarily in a global reduction of volatile production (Raguso *et al*., 2007; Doubleday *et al*., 2013; Sas *et al*., 2016; Majetic *et al*., 2019; Jantzen *et al*., 2019; Petrén *et al*., 2021). It remains unclear, however, whether independent transitions to selfing lead to similar changes in floral fragrance.

The genus *Capsella* provides a genetically tractable model to study the basis of the repeated evolution of selfing syndrome traits (Wozniak *et al*., 2020). In this genus, the transition to selfing occurred at least two times independently through the breakdown of the self-incompatibility system of the outcrossing *C. grandiflora* (*Cg*)-like ancestors. One of these events occurred about ∼200kya, giving rise to *C. rubella* (*Cr*) (Foxe *et al*., 2009; Guo *et al*., 2009; Koenig *et al*., 2018). The timing of the emergence of the second selfing lineage, *C. orientalis* (*Co*) is less clear but is likely to have occurred within the last 2 million years (Hurka *et al*., 2012; Bachmann *et al*., 2018). In contrast to *C. rubella* and *C. orientalis*, the outcrossing species *C. grandiflora* emits high levels of benzaldehyde, indicating that the composition of flower fragrance has changed in both selfers (Sas *et al*., 2016; Jantzen *et al*., 2019). The extent to which the evolution of flower scent in *Cr* and *Co* has been similar is, however, not known.

In this study, we took advantage of the *Capsella* genus to investigate the convergent evolution of floral fragrance after independent transitions to selfing. While the amount of volatiles emitted did not decrease in the two selfers, both show similar changes in floral scent composition. The most common change was the reduction of (*E*)-*β-*ocimene production. We demonstrate that this was achieved through different genetic mechanisms. In *C. rubella*, it is linked to a non-synonymous mutation in the chloroplast transit signal of the homolog of the *A. thaliana TERPENE SYNTHASE 2* (*TPS02*) that seems to decrease the efficiency of chloroplast targeting. Population genetic analyses support a scenario in which this mutation has been fixed in the selfing *C. rubella* lineage by capturing an allele segregating in the ancestral outcrosser species. Our results suggest that certain volatile compounds are more likely to be lost or gained after the transition to selfing. The fact that the reduction of (*E*)-*β-*ocimene emission evolves from the fixation of a rare variant in the ancestral standing genetic variation may suggest that it provides a selective advantage at the early step of selfing evolution.

## Results

### Independent transitions to selfing in the genus Capsella are accompanied by convergent changes in floral fragrance

To determine to which extent independent events of selfing evolution are associated with similar changes in flower volatile emission, we analyzed the floral fragrance of 5 representative accessions for the outcrossing species *C. grandiflora* (*Cg*) and its two incipient selfers, *C. rubella* (*Cr*) and *C. orientalis* (*Co*). Historically, *Cg* has split into an eastern and western lineage from which *Cr* and *Co* have evolved, respectively (Hurka *et al*., 2012). Modern *Cg* descend from the western lineage while ‘eastern’ outcrossing populations are believed to be extinct. Consistently, hierarchical clustering based on genetic distances, using available genomic datasets, places *Cr* in a sister clade to *Cg* while *Co* branches as an outgroup (**Figure 1A**).

**Figure 1:**
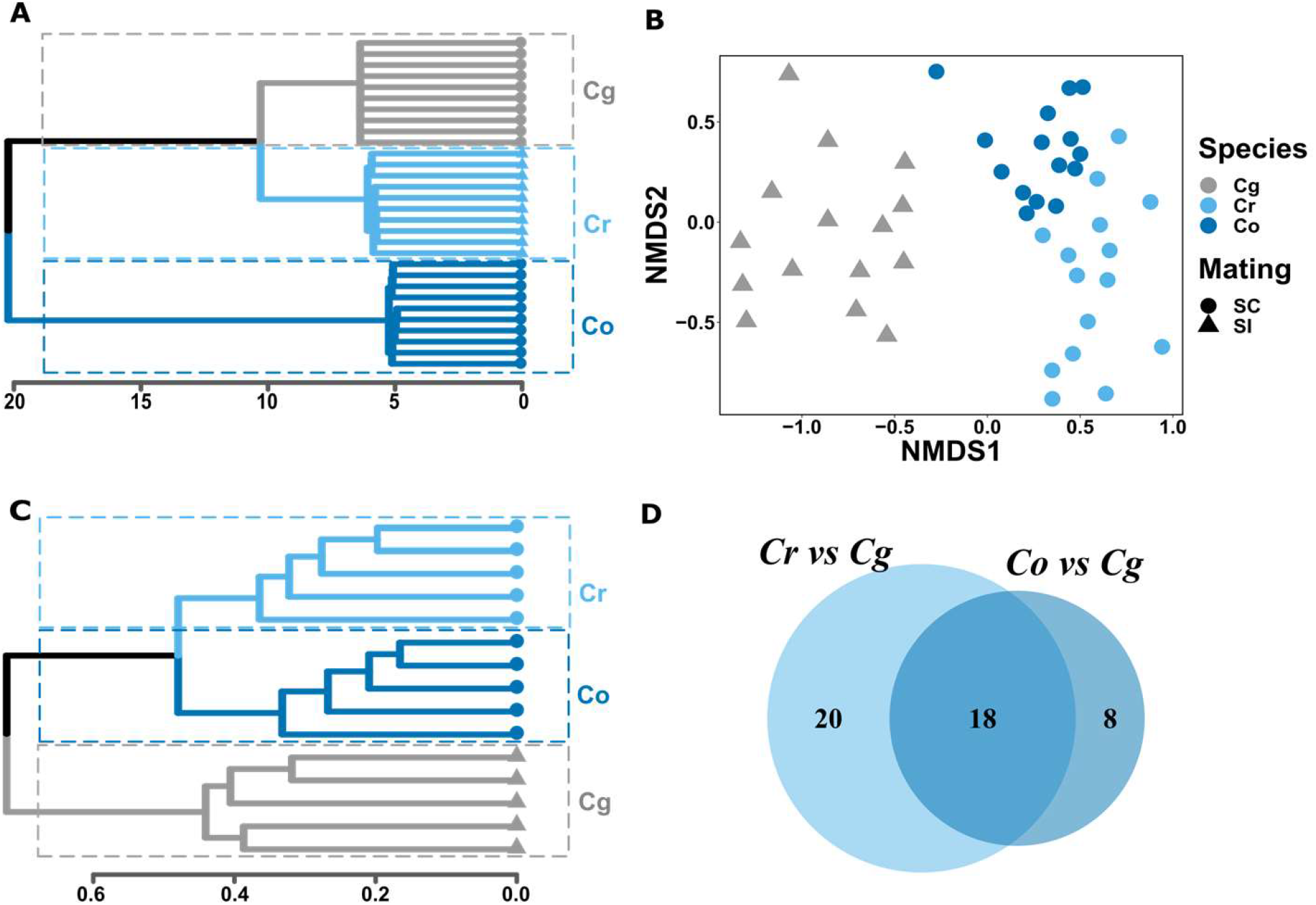
Convergent evolution of flower scent after independent transitions to selfing in the genus *Capsella*. **(A)** Hierarchical clustering using Ward clustering algorithm on hamming distances between 10 *Cg, Cr* and *Co* populations. The scale bar indicates log(hamming distances). (**B**) Non-metric multidimensional scaling (NMSD) plot based on Bray-Curtis dissimilarities in flower scents composition between samples (n =15 per species). (**C**) Clustering cladogram illustrating the similarities between the flower scent composition of all samples tested based on average Bray-Curtis dissimilarities. The scale bar indicates Bray-Curtis dissimilarities. (**D**) Venn diagram illustrating the number of compounds significantly changed only in *Cr*, only in *Co* or in both *Co* and *Cr* compared to *Cg*. SI and SC refer to self-incompatible and self-compatible, respectively.

Our analyses detected 142 volatile compounds over all samples. Total flower emission did not differ significantly between mating types; only *Co* emitted a slightly higher amount of volatiles (**Figure S1**). Nevertheless, we observed extensive variation in scent composition. Non-metric multidimensional scaling (NMDS) plots of dissimilarity indices, calculated based on the proportion of all the volatile compounds consistently detected in at least one species, differentiate the individuals according to their mating types (**Figure 1B and C**). While samples from different species cluster together, individuals from both *Co* and *Cr* displayed higher NMDS1 values compared to outcrosser individuals (**Figure S1**). Contrary to what could be expected from genetic distances (**Figure 1A**), clustering of the different samples based on the flower scent composition grouped *Cr* and *Co* as sister clades while *Cg* branched as an outgroup (**Figure 1C**). Such clustering indicates that selfers’ flower fragrances were more closely related to each other than to those of the outcrossers. Indeed, the mating system explains most (about 40%) of the variation in flower scent composition (PERMANOVA - *F* = 43.128, *p* < 0.001), while species and population explained only 11 and 22 %, respectively (**Table S2**). We next performed a Random forest analysis to examine the extent to which different populations could be classified into different mating groups based on the flower volatile emission profile. The model built on the relative proportions of volatile compounds correctly classified all populations with a probability to be assigned to the right category above 75% (**Figure S2A and B**). To determine the importance of different volatile compounds for this classification, we calculated the mean decrease in model accuracy when the different volatiles are removed from the dataset (**Figure S2C**). Nineteen compounds are particularly important to assign the different populations to the correct mating type categories, with (*E*)-*β*-ocimene, (*Z*)-*β*-ocimene and benzenacetaldehyde causing the largest drop in classification accuracy. Overall, 38 and 26 compounds vary significantly between *Cg* and *Cr* or *Cg* and *Co*, respectively (**Figure 2 and S3, Table S3 and S4**). Among those, we found a significant overlap of 18 compounds (*p* = 0.01 – hypergeometric probability) which comprise almost all of the ninteen compounds important for the classification into different mating classes (**Figure 1D**). All these compounds change in the same direction in both selfers but not always to the same degree. Seven of these compounds were not different between both selfers, while the remaining four also differed significantly between *Co* and *Cr* (**Figure S3**). Globally, *Cr* tends to have greater changes in volatile emissions, suggesting that the molecular process underlying these changes may differ between the two selfing lineages. Volatile emissions were not always lost in the selfers as two compounds were also found to be emitted at a higher level in selfing species (**Figure S3**)

**Figure 2:**
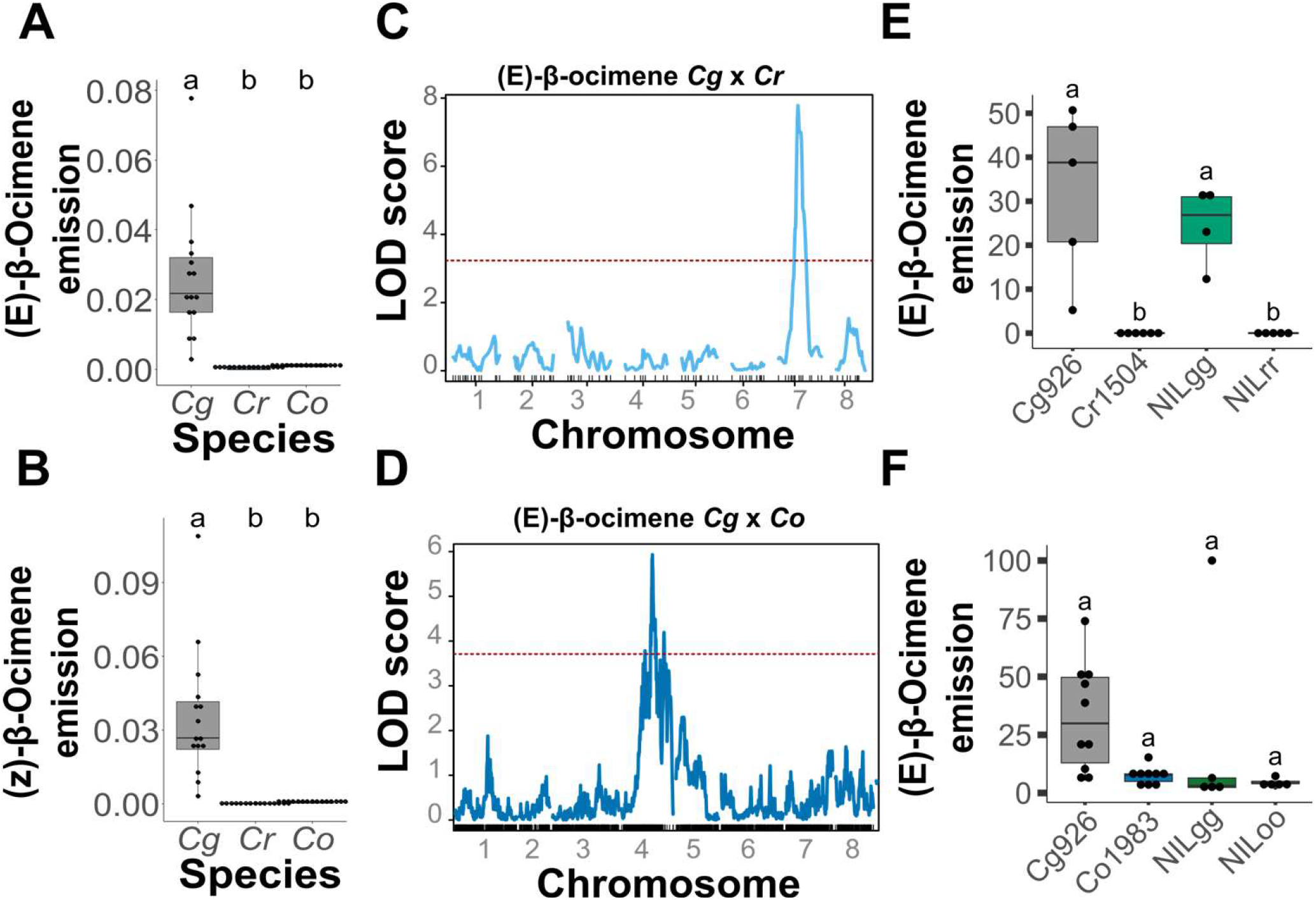
Mutations at different loci led to the loss of β-ocimene in *C. rubella* and *C. orientalis*. (**A**) and (**B**). The relative abundance of (*E*)-*β*-ocimene and (*Z*)-*β*-ocimene in each *Capsella* species (n=15 for each species). (**C**) and (**D**). QTL mapping for the emission of (*E*)-*β*-ocimene (Tag_mass 93) in the *Cr1504* x *Cg926* RIL population and (**C**) *Co1983* x *Cg926* F2 population (D). Red dashed lines indicate the genome-wide 5% significance threshold. (**E**) and (**F**) Quantification of (*E*)-*β*-ocimene emission (Tag_mass 93) in the headspace of NILs segregating for the *Cg* and *Cr* alleles (**E**) or the *Cg* and *Co* alleles (**F)** (n=5). The intensity of the (*E*)-*β*-ocimene detected for each individual was scaled to maximal intensity in the experiment and presented as percentage value. Letters indicate significant differences as determined by one-way ANOVA with Tukey’s HSD test (α=0.05).

These results indicated a convergent evolution of volatile production after independent transitions to selfing in the genus *Capsella*. The similarities are such that it is possible to classify individuals into different mating types solely based on the floral fragrance composition. The fact that the emission of some volatile compounds increased in the two selfing lineages further suggests that the changes in flower scent emission are not simply a result of resource allocation, but could also reflect an ecological adaptation to a new mode of reproduction.

### Mutations at different loci underlie the loss of (E)-β-ocimene emission after the transition to selfing in Capsella

The convergent evolution of flower scent observed between *Cr* and *Co* offers the opportunity to investigate the genetic basis of flower scent evolution and to determine whether independent changes in the emission of the same volatile compounds tend to involve mutations within the same genes. We, therefore, mapped Quantitative Trait Loci (QTL) influencing the emission of the most differentiated compounds between mating categories, (*E*)-*β-*ocimene and benzeneacetaldehyde in both *Cr* and *Co*.

(*E*)-*β-*ocimene and (*Z*)-*β-*ocimene were detected in all *Cg* samples but were not emitted by any of the *Cr* individuals analyzed and produced at very low levels in *Co* (**Figure 2 A and B**). Since both compounds are produced through the same biosynthetic pathway, with a bias towards the production of the (*E*) stereoisomer in many species (Farré-Armengol *et al*., 2017), we focused further analyses on (*E*)-*β*-ocimene. Only 50% of the *Cr* x *Cg* RILs emitted (*E*)-*β-*ocimene to a detectable amount corresponding to the expected segregation ratio caused by mutations at a single locus (**Figure S4**). Accordingly, we detected only one major QTL (QTL_1), located on chromosome 7 and explaining about 51% of the total variance in (*E*)-*β-*ocimene emission in this population (**Figure 2 C; Table S5**). In the *Co* x *Cg* F2 population, (*E*)-*β-*ocimene was detected in 99.4% of the samples, with a distribution of volatile emission skewed toward small values (**Figure S4B**). The analysis of the segregation ratio in this population suggested that at least two loci could contribute to the loss of (*E*)-*β-*ocimene in *Co* (**Figure S4F**). Nevertheless, our QTL analysis detected only one QTL (QTL_2) located on chromosome 4, explaining only 7.6% of the total variance (**Figure 2 D; Table S5**). Similarly, the loss of benzeneacethaldehyde emission appears to be controlled by two loci in *Cr* and a single locus in *Co*, in which additional loci likely influence its emission levels quantitatively. Our genetic analysis revealed a QTL located on chromosome 3 in the *Cr* x *Cg* RILs, but we did not detect any significant QTL in the *Co* x *Cg* F2 population. Together, these results suggest a different genetic basis for the losses of the emission of these two compounds in *Co* and *Cr*. It is nevertheless important to note that our QTL model poorly explains the phenotypic variation in our dataset, particularly in *Co*, most likely because of a highly complex genetic basis and/or an important non-genetic variance limiting the statistical power of our analysis.

To ascertain that different loci underlie the loss of scent emission in the two *Capsella* selfing species, we formally tested whether allele variation in the QTL_1 influence (*E*)-*β-*ocimene emission in both selfers. To this end, we generated Near Isogenic Lines (NIL) segregating for either the *Cr* and *Cg* alleles (*NILrg)* or the *Co* and *Cg* alleles (*NILog*) in a chromosomal region spanning QTL_1 but otherwise homozygous for the selfer alleles throughout most of the genome. We next selected plants homozygous for alternative alleles and compared their scent emission profile (**Figure 2 E and F**). The *NILrg* individuals homozygous for the *Cg* allele (*NILgg1*) emit (*E*)-*β-*ocimene to a similar level as wild-type *Cg* plants. As for *Cr*, no (*E*)-*β-*ocimene could be detected in *NILrg* homozygous for the *Cr* allele (*NILrr*). This result confirmed our ability to detect QTL influencing volatile emission and further demonstrated that allelic variation at this locus can fully explain the loss of (*E*)-*β-*ocimene emission in *Cr* (**Figure 2 E**). By contrast, both plants homozygous for the *Cg* allele (*NILgg*) and those homozygous for *Co* (*NILoo*) emitted (*E*)-*β-* ocimene at a detectable and similar level, further suggesting that QTL_1 does not influence (*E*)-*β-* ocimene production in *Co* (**Figure 2 F**). Therefore, at least in the case of (*E*)-*β-*ocimene, the loss of scent is not caused by mutation at the same loci.

### Capsella TPS02 underlies the loss of (E)-β-ocimene in C. rubella

To better understand the history and molecular basis of flower fragrance evolution after the transition to selfing, we set ourselves to identify the gene underlying the loss of *(E)-β-*ocimene in *Cr*. To this end, we screened about 1,200 progeny plants of *NILrg* for plants harbouring a recombination event within the QTL_1 region. Comparing the level of *(E)-β-*ocimene production with the position of the recombination breakpoints allowed us to narrow down the casual mutations to a region of 115 kb comprised between 9.330.402-9.445.345Mb on chromosome 7 (**Figure 3A**). While all individuals heterozygous within this interval emitted a detectable amount of *(E)-β-* ocimene (average levels of 3 to 15% of the maximum intensity of (*E*)-*β*-ocimene detected), this compound could not be detected in any individuals homozygous for the *Cr* allele. Within this region, we identified two genes coding for terpene synthases (TPS) in a tandem repeat. These genes were syntenic to *Arabidopsis thaliana TPS02* and *TPS03*, with an average protein sequence similarity of 85% and 84% (**Figures 3B and S5**). Both genes share a similar structure, with seven exons and six introns, and a high level of sequence conservation between the two paralogues (53.6% and 52.8% in *Cg* and *Cr*, respectively), but TPS02 is 35 (36 in TPS02g) amino acids longer than TPS03 (**Figure S5**). TPS02 has previously been shown to encode (*E*)-*β*-ocimene synthase in *A. thaliana* (Huang *et al*., 2010). It was further demonstrated that the presence of a chloroplast transit peptide (cTP) is necessary for the proper TPS02 subcellular localisation and synthesis of (*E*)-*β*-ocimene (Huang *et al*., 2010). In silico prediction of target peptides identified a cTP in both *Cr* TPS02 and *Cg* TPS02 but no target peptide was identified in any of the *TPS03* alleles (**Table S6**). Accordingly, we observed that TPS02 is targeted to the chloroplast in *Cg* (**Figure 4B and C**), suggesting that it may also contribute to (*E*)-*β*-ocimene in *Capsella*. Based on these observations, we hypothesized that allelic variation at TPS02 may underlie the loss of (*E*)-*β*-ocimene emission in *Cr*. To test this hypothesis, we compared the ability of genomic fragments encoding for *Cg TPS02* (TPS02g) or *Cr TPS02* (TPS02r) to restore *(E)-β-*ocimene in the *NILrr*. While *(E)-β-*ocimene could not be detected in any of the 10 *TPS02r* lines tested, all the lines transformed with *TPS02g* emitted a measurable amount of this compound (**Figure 3C**). Together, these results indicated that allelic variation between *Cr* and *Cg* are responsible for the change in *(E)-β-*ocimene emission between these species and that introducing *CgTPS02* into *Cr* was sufficient to fully rescue the emission of this compound.

**Figure 3:**
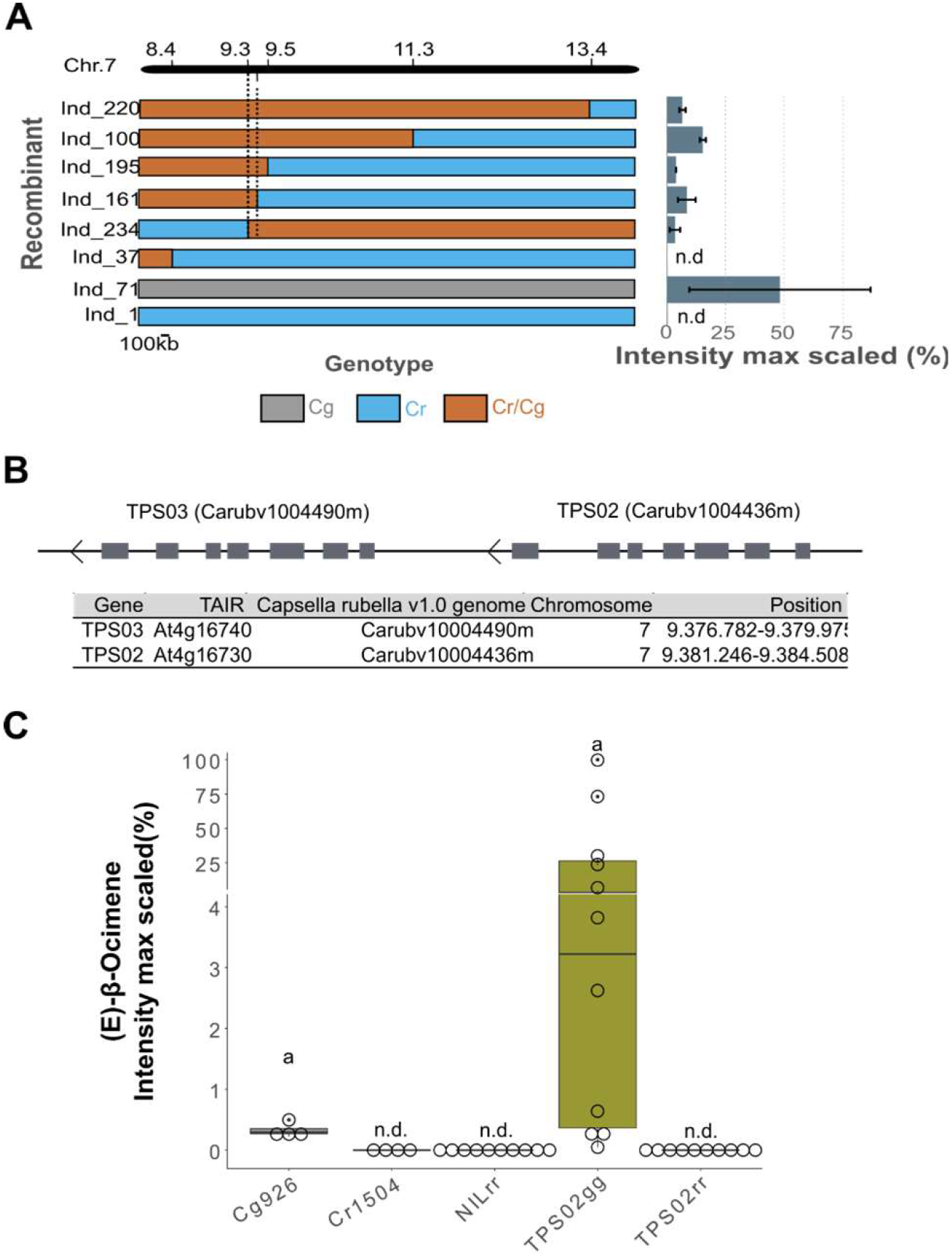
The loss of *β*-ocimene emission in *C. rubella* maps to TPS02. (**A**) Fine-mapping of the QTL_1 controlling (*E*)-*β*-ocimene emission. Bars on the left indicate the genotype of informative recombinants. Positions on chromosome 7 are indicated in Mb on the top, and the genotypes of the different recombinants along the region of interest are color coded. The bar chart on the right indicates (*E*)-*β*-ocimene emission intensity scaled to the maximal value in the experiment for each recombinant. Values are mean ± SD from 2 measurements. (**B**) Schematic representation and genomic locations of genes known to be involved in monoterpene biosynthesis and present within the mapping interval. (**C**) (*E*)-*β*-ocimene emission intensity scaled to the maximal value in *Cg926, Cr1504, NILrr* and *NILrr* transformed with either the *Cg TPS02* (*TPS02g*) or the *Cr TPS02* (*TPS02r*) alleles. Each dot represents an individual measurement. Letters indicate significant differences as determined by one-way ANOVA with Tukey’s HSD test (α=0.05).

**Figure 4:**
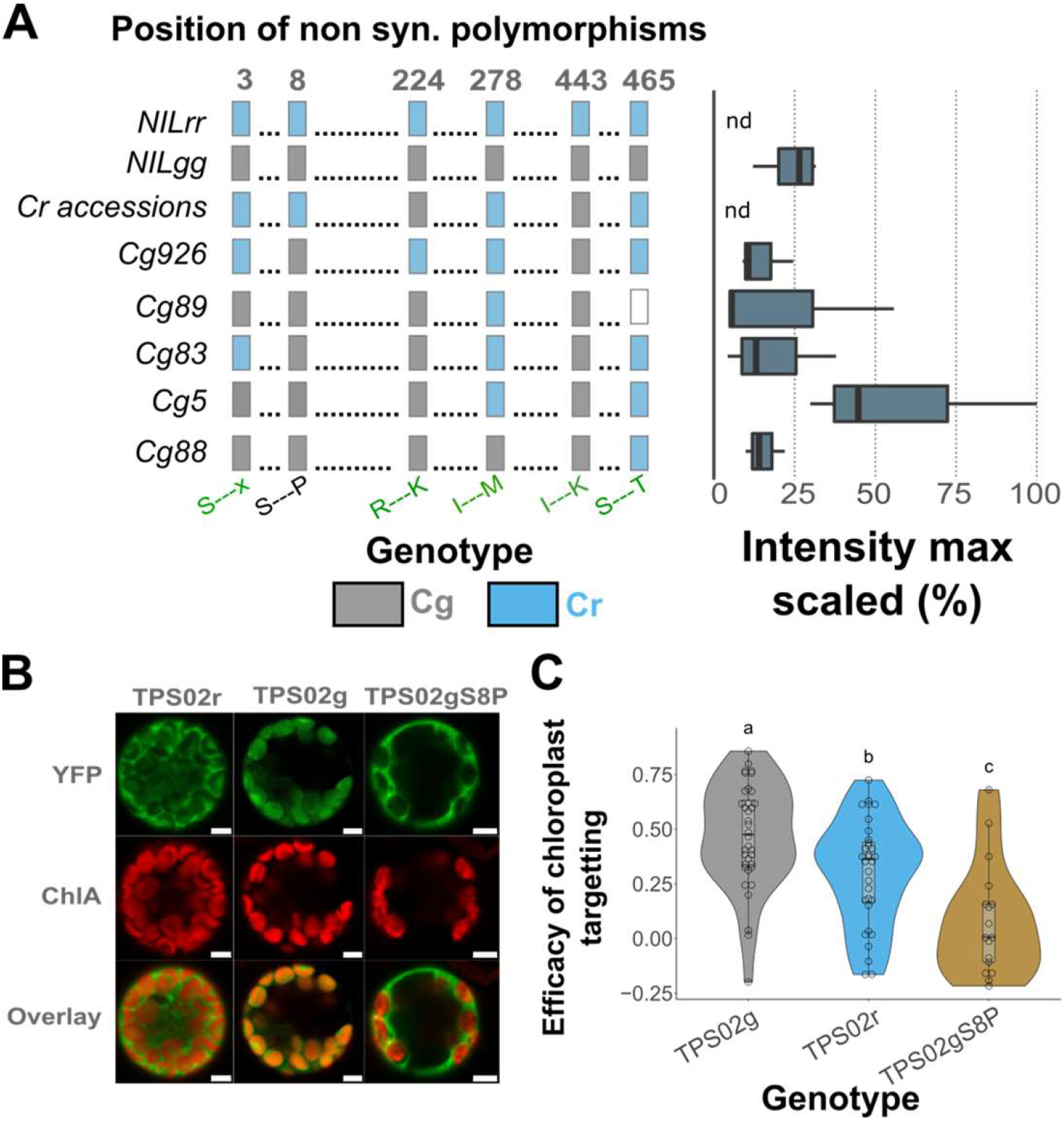
A mutation in the chloroplast transit peptide sequence underlies the loss of *β*-ocimene emission in *C. rubella*. (**A**) Comparison between (*E*)-*β-*ocimene (Tag_mass 93) emission and the genotype at the 6 candidate nonsynonymous polymorphisms in different *Cg* and *Cr* accessions. The genotype at each polymorphic amino acid position is represented on the right by color coded rectangles. Boxplots on the right indicate the intensity of the (*E*)-*β -*ocimene emitted by each individuals scaled to the maximal value in the experiment. (**B**) Representative images of the cellular localization of CrTPS02 (TPS02r), CgTPS02 (TPS02g) or CgTPS02 in which the serine at position 8 has been converted into a proline (TPS02gS8P) in *Arabidopsis* leaf protoplasts. YFP corresponds to the cellular localization of TPS02, the Chlorophyll A signal (ChlA) indicates the position of the chloroplasts and the overlay illustrates the overlap between the two signals. Scale bars correspond to 5 μm. (**C**) Violin plot representing the efficiency of TPS02r, TPS02g and TPS02gS8P chloroplast targeting estimated as the above threshold Pearson correlation coefficient between the YFP and ChlA signals. Each dot represents an individual measurement.

### Inefficient targeting of TPS02 to the chloroplast underlies the loss of (E)-β-ocimene in C. rubella

To get insights into the molecular mechanism underlying the loss of *(E)-β-*ocimene emission, we first compared *TPS02* expression level and mRNA splicing pattern between *NILgg* and *NILrr* plants (**Figure S6**). *TPS02* was expressed at similar levels in the inflorescences of both genotypes, which also transcribed an mRNA of similar size and structure (**Figures S5 and S6**). This lack of qualitative and quantitative differences in *TPS02* transcription suggests that the loss of (*E*)-*β*-ocimene emission has been caused by mutations affecting the TPS02 amino acid sequence. The comparison between TPS02 alleles segregating in our mapping population (i.e., a cross between *Cr1504* and *Cg926*) identified six non-synonymous mutations, including five amino acid exchanges and one nucleotide triplet deletion (**Figure S5**). To pinpoint which of these amino acids may be responsible for the loss of *(E)-β-*ocimene emission, we exploited the large genetic diversity of *Cg*. We correlated the genotype at the candidate non-synonymous polymorphisms with *(E)-β-* ocimene emission in five *Cg* and five *Cr* plants from different accessions (**Figure 4A**). *(E)-β-* ocimene was emitted at various levels of all *Cg* plants analyzed but was not detected in any *Cr* accessions analyzed. For all but two non-synonymous SNPs (at position 9.382.389/ amino acid 443 and 9.384.490/ amino acid 8), we found at least one *Cg* individual homozygous for the *Cr* allele, indicating that these polymorphisms were not causal. One of the two remaining polymorphic sites (position 9.382.389/ amino acid 443) was, however, homozygous for the *Cg* allele in all tested *Cr*, excluding it from the list of plausible causal mutations (**Figure 4A**). Only the polymorphism at position 9.384.490 (8^th^ amino acid) fully correlates with the production of (*E*)-*β*-ocimene in our assay, making it the prime candidate to underlie the loss of *(E*)-*β*-ocimene.

The corresponding SNP induces a serine-to-proline conversion, which corresponds to a polar to non-polar amino acid exchange within the cTP (**Figure S5**). Because TPS02 chloroplast localization is essential to *(E)-β-*ocimene synthesis (Huang *et al*., 2010), we sought to test if changes in its cellular localization may be the basis of the loss of (*E*)-*β*-ocimene emission. To assess the cellular localization of TPS02r and TPS02g, we introduced a yellow fluorescent protein (YFP) before their Coding DNA Sequence (CDS) stop codon. We transiently expressed these recombinant proteins in *A. thaliana* protoplasts and estimated the efficiency of chloroplast targeting by determining the above threshold correlation coefficient between the YFP and the Chlorophyll A (present within the chloroplast) fluorescent signals (**Figure 4B and C**). This analysis indicated that a significantly larger proportion of the TPS02g is localized within the chloroplast, suggesting that allelic variation between these two proteins affects their cellular localization. We thus mutated nucleotide 9.384.490 in TPS02g to introduce a proline while keeping the rest of TPS02g protein unchanged. This point mutation decreased the efficiency of TPS02g targetting to the chloroplast, suggesting that the mutation at SNP 9.384.490 affected the function of the Chloroplast transit signal and reduced the amount of TPS02 in the chloroplast (**Figure 4B and C**). This change has, therefore, likely impaired the conversion of geranyl diphosphate to (*E*)-*β*-ocimene compound inside the chloroplast, reducing this overall emission.

### Evolutionary history of the loss of (E)-β-ocimene

To gain insights into the evolutionary origin of the loss of *(E)-β-ocimene* emission, we further investigated the genetic diversity at *TPS02* within *Capsella*. We reconstituted the TPS02 haplotypes from 182 *Cg*, 16 *Co* and 51 *Cr* accessions, focusing mainly on their CDS. We first investigated the genetic structure at the *TPS02* locus within the *Cr* dataset by inferring individuals’ genetic ancestries using ADMIXTURE (**Figures 5A and S7**). The best fit of individuals’ ancestries was obtained with six clusters, with two of these clusters including 37 out of 51 individuals. Hierarchical clustering of the *Cr* haplotypes identified two major clades regrouping together 48 *Cr* individuals, later referred to as the Crub_183 and Cr1504 haplotype groups. The geographical distribution of these haplotypes does not reflect the genetic relationship between *Cr* individuals suggesting that they were differentiated at the early stage of *Cr* history before its geographical spread (**Figure 5A**). To understand how these different clusters relate to the *TPS02g* haplotypes, we reconstituted a Neighbor-Joining tree using all *TPS02* assembled haplotypes (**Figure 5B**). The haplotypes corresponding to the 48 individuals mentioned above cluster within a lineage also containing haplotypes found in 8 *Cg* individuals. Nevertheless, the latter show excessively long *Cr*-like haplotypes and low genetic differentiation from *Cr*-haplotypes (**Figure 5C and S8**). Such genomic signature is reminiscent of recent hybridization events indicating that these 11 *Cr*-like haplotypes have been recently introgressed from *Cr* into *Cg* and have not yet been broken up by recombination events. The remaining *Cr* haplotypes (corresponding to three individuals) cluster within *Cg* haplotypes. In addition to unusual genetic similarities to *Cg*, these haplotypes show particularly low levels of long *Crub_183* or *Cr1504* haplotypes (Figure S8). Furthermore, the individuals harbouring these haplotypes all originate from Greece where the two *Capsella* species remain in sympatry. Together with the unusual polymorphism pattern, it suggests that these haplotypes correspond to post-divergence gene flow from *Cg* to *Cr*.

**Figure 5:**
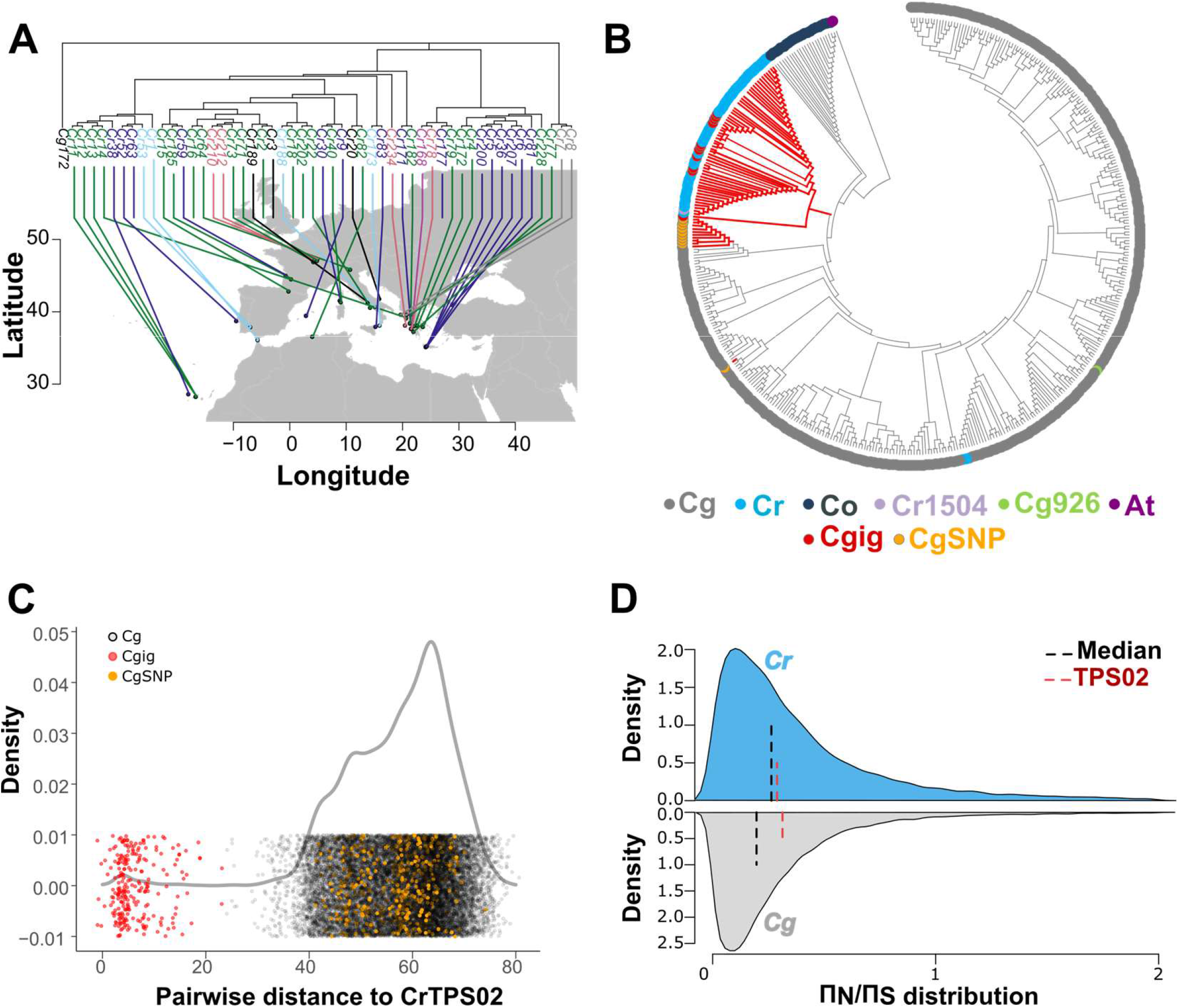
Evolutionary history of TPS02 in *Capsella*. (**A**) Geographical distribution of the different *Cr* TPS02 haplotype groups, as determined by an admixture analysis (**Figure S6**). The tree on the top illustrates the phylogenetic relationship among corresponding *Cr* accessions determined based on genome-wide SNPs information. (**B**) Neighbor-Joining tree representing the genetic relationships between all *Capsella* TPS02 haplotypes identified. The bootstrap consensus tree was inferred from 1000 replicates. Branches corresponding to partitions reproduced in less than 50% bootstrap replicates are collapsed. Red branches illustrate haplotypes containing the candidate causal mutation at SNP 9.384.490. *Cg* haplotypes containing the *Cr* allele at this position are marked by a red dot if they show signs of recent introgression (Cgig) or by an orange dot if they do not show any sign of recent introgression. (CgSNP). (**C**) Distribution of the pairwise genetic distances, as numbers of polymorphisms, between *Cg* and *Cr* TPS02 haplotypes. Values corresponding to distance between *Cgig* and *Cr* haplotype are shown in red, while the values comparing *CgSNP* and *Cr* haplotypes are shown in orange. (**D**) Π_N_/Π_S_ distribution in *Cg* and *Cr*. Median and TPS02 Π_N_/Π_S_ are indicated with a black and red dashed line, respectively.

After excluding individuals with genomic signatures of recent introgression, we observed that the *Cr* rubella allele at position 9.384.490, causing the serine-to-proline conversion, was found in all *Cr* but also in 11 *Cg* individuals (**Figures 5 and S8**). None of these *Cg* individuals shows signs of recent introgression. Their pairwise distances to *Cr* haplotype are dispersed through the centre of the distribution of all *Cg* haplotypes, and they show average levels of long *Cr*-like haplotype. All non-synonymous mutations within *TPS02*, were found to similarly segregate in *Cg* (**Figure S8**). Together these observations support a scenario in which the *Cr* allele leading to the loss of (*E*)-*β*-ocimene has been captured from standing variation in the ancestral population. Selfing syndrome traits such as scent emission are often believed to evolve as a result of the relaxation of selective pressure acting on genes controlling pollinator attraction. To determine if the functional constraints acting on TPS02 have indeed been relaxed in *Cr*, we computed the ratio between non-synonymous sites and synonymous sites’ nucleotide polymorphism (Π_N_/Π_S_). Consistently with the expectation of the decreased efficiency of natural selection in selfers (Slotte et al. 2013), the genome-wide Π_N_/Π_S_ distribution shifted towards higher values in *Cr* (**Figure 5D**). However, TPS02 Π_N_/Π_S_ did not change between the two species, suggesting that similar levels of functional constraints are still driving the evolution of this gene in *Cr*. This data suggests, therefore, that the loss of (*E*)-*β*-ocimene emission in *Cr* does not seem to have been caused by the relaxation of purifying selection acting on the gene controlling its production.

## Discussion

In the *Capsella* genus, the transitions to selfing did not lead to global changes in the total amount of volatiles emitted but rather affected the production of specific compounds. The similarity between the flower scent evolution in the two selfers is such that its composition can accurately predict the mating systems. In total, we found 18 compounds similarly affected in both selfers. The emission of 16 of them was reduced after the switch in the mating system, which is consistent with a loss of pollinator’s attractive features triggered by the evolution of self-fertilization. Among them, we were able to annotate five of these compounds, including (*E*)-*β*-ocimene, (*Z*)-*β*-ocimene, benzeneacetaldehyde, which are common volatiles in floral bouquets believed to promote pollinator attraction (Knudsen and Tollsten, 1993; Knudsen *et al*., 1993, 2004, 2006; Knudsen and Gershenzon, 2006; Weiss *et al*., 2016; Qian *et al*., 2019). In particular, *β*-ocimene has been proposed to act as a generalist pollinator attractant since it is emitted by over 50% of angiosperm families, and it has been directly shown to increase the rate of pollinator visitations (ANDERSSON *et al*., 2002; Filella *et al*., 2013; Byers *et al*., 2014; Farré-Armengol *et al*., 2017, 2020; Peng *et al*., 2017). Nevertheless, our analysis also revealed that compounds involved in defence mechanisms are also affected by the transition to selfing. Allyl isothiocyanate is a toxic compound also acting as a repellent against various organisms, including bacteria, fungi and herbivores (Agrawal and Kurashige, 2003; Clay *et al*., 2009; Müller *et al*., 2018; Yang *et al*., 2021; Singh *et al*., 2022). Strikingly, its emission is also reduced in the two selfers suggesting that the reproduction optimization in animal pollinated plants relies on a delicate balance between attraction and defence mechanism. Outcrossing plants likely emitted a complex mixture of volatiles that promote pollinators’ visitations while limiting damage from insect pests and their associated pathogens (Jacobsen and Raguso, 2018). Therefore, it is plausible that because the decrease in pollinator attractiveness also reduces the attraction of pests, the transition to selfing relieves the requirement for strong defence mechanisms against herbivores. We also observed that the emission of two compounds increased in both selfers. One of them was annotated as dimethyl trisulfide, which has an unpleasant sulfurous odor known to be attractive to silphid beetles and some dipterans (Podskalská *et al*., 2009; Zito *et al*., 2014; von Hoermann *et al*., 2016; Yan *et al*., 2018). Although it remains to be determined whether the increased emission of such compounds presents a fitness advantage to self-fertilizing plants or simply reflects metabolic trade-offs, it suggests that energy is still invested in producing flower signals in self-fertilizing plants.

The convergent evolution of flower scent in both selfing lineages is not caused by mutations within the same genes. Our results indicate that the reduction of both *β*-ocimene and benzeneacetaldehyde have a different genetic basis in *Cr* and *Co. β*-ocimene emission was not detected in *Cr*, while it could be detected at lower levels in some of the *Co* accessions tested. Accordingly, QTL influencing its emission mapped to different genomic locations in both selfers. In *Cr*, the loss of *β*-ocimene was caused by a mutation affecting *TPS02*, but allelic variation in *TPS02* between *Co* and *Cg* did not significantly change this compound’s emission. The most probable causal SNP is located within a chloroplast transit signal of TPS02 and appears to impair its chloroplastic localization. Monoterpenes, such as (*E*)-*β-*ocimene are synthesized in plastid compartments through the methylerythritol 4-phosphate pathway (MEP) (Pichersky *et al*., 1994; Guterman *et al*., 2002; Chen, 2003; Tholl *et al*., 2005; Nagegowda *et al*., 2008). Their productions, therefore, relies on the localization of monoterpene synthase within the plastids (Turner *et al*., 1999). In contrast, sesquiterpenes are produced in the cytosol via the mevalonic acid pathway (MVA) (Dudareva and Pichersky, 2000; Tholl and Lee, 2011). In most plants, there is limited crosstalk between both compartments, and the importance of the plastidic localization of monoterpene synthases has been reported for a few of them, including TPS02 (Bouvier *et al*., 2000; Nagegowda *et al*., 2008; Huang *et al*., 2010). Although our candidate mutation did not totally impair TPS02 chloroplastic localization, it is plausible that the decrease in the amount of chloroplastic TPS02 limits *β-*ocimene biosynthesis, preventing its detection within the floral blend. The fact that TPS02 has been shown to be capable of synthesizing the sesquiterpene (*E,E*)-*α*-farnesene in *A. thaliana* (Huang *et al*., 2010), suggests that the cytosolic TPS02 now contribute to the mevalonic acid pathway (MVA) in *Cr*. As such, the causal mutation is likely to have minimal economic implications by rechannelling the enzyme towards other metabolic pathways. Although (*E, E*)-*α*-farnesene has been detected in the floral blend of several angiosperm species, its emission has most often been found to be activated upon herbivore attack as an indirect defence mechanism attracting natural pests’ enemies (Boevé *et al*., 1996; Scutareanu *et al*., 2003; Danner *et al*., 2011; Lin *et al*., 2017). Based on this, it could be hypothesized that changes in TPS02 subcellular localization allow shifting the floral blend’s function from pollinator attraction to defence mechanism. The QTL influencing *β-*ocimene emission in *Co* does not contain any known *β-* ocimene synthase indicating that a different type of mechanism is involved. It is, therefore, unclear whether the decrease in *β-*ocimene production may also favour other metabolic pathways and ecological functions in *Co*. The fact that different loci have caused the reduction of *β-*ocimene emission in both selfers suggests that no strong genetic constraints influence the evolution of floral volatiles. Therefore, the convergent evolution of floral scent is more likely only to reflect the changes in ecological pressures associated with the shift in the mating system.

So why do independent transitions to selfing affect the emission of the same volatiles? The fact that similar compounds are affected in both of the selfing lineages indicates that these compounds are particularly advantageous in outcrossers, while they do not provide fitness advantages in self-fertilizing plants. The lack of changes in the extent of functional constraint acting on *TPS02* argues against a general release of purifying selection on the genes controlling *β*-ocimene production in selfers. Their emission could in principle be reduced because of high energetic costs or fitness penalties it incurs to selfing lineages. Although the carbon allocation to volatile emission is likely minor compared to changes in flower morphologies (Grison-Pigé *et al*., 2001), the production of volatiles, such as terpenes, is believed to be energetically costly (Gershenzon, 1994). Yet, total volatile production is not affected in *Capsella* selfers. Furthermore, it is unlikely that the loss of specific compounds, like *β*-ocimene, provides large energetic benefits, especially in *Cr*. Indeed, our data suggest that a non-synonymous mutation in *TPS02*, an enzyme involved in a terminal step of monoterpene production, caused the loss of *β*-ocimene in *Cr*. In addition, this mutation did not affect TPS02 metabolic activity but rather its subcellular localization. This protein may thus still contribute to the production of other sesquiterpenes in *Cr*, making any energetic gain associated with the underlying mutation likely minor. Moreover, the fact that all *Cr* accessions shared this mutation, which can still be found in *Cg*, indicates that the loss of *β*-ocimene emission evolved early after the emergence of self-fertilization, most likely through the capture of an allele segregating in the outcrossing ancestors. Based on these observations, it is tempting to speculate that reducing the emission of volatiles attractive to pollinators provides an advantage at the early stages of selfing evolution. Self-compatible genotypes less attractive to pollinators could have a higher chance of establishing selfing lineages as the associated reduction in conspecific pollen flow would increase the probability of self-fertilization. Testing this hypothesis will, nevertheless, require quantifying the energetic and reproductive benefits of the loss of individual compounds in early emerging selfing lineages.

The convergent evolution of flower scent, together with the evolutionary history and low pleiotropy of the mutation underlying the loss of *β*-ocimene emission in *Cr*, suggest that the evolution of flower chemical signals in selfers cannot simply be explained by the relaxation of purifying selection in outcrosser lineages. The floral bouquet’s final composition is likely to be guided by a shift in the ecological pressures to maintain or inhibit pollinator attraction and defence mechanisms. Dissecting the biological functions of convergent selfing syndrome traits should therefore help to understand the evolutionary and ecological pressures shaping flower phenotypes and thus identify important factors in the optimization of different modes of plant reproduction.

## Materials and Methods

### Biological Materials

The *Capsella grandiflora* (*Cg*), *C. rubella* (*Cr*) and *C. orientalis* (*Co*) accessions used in this study had been collected from diverse locations in Europe, Africa, and Asia. Detailed information on the locations is given in **Table S1**. The *Cr* x *Cg* RILs and *Co* x *Cr* F2 population used for the QTL mapping experiments have been previously described and were generated from crosses between *Cr1504* and *Cg926* and *Co1983* and *Cg926*, respectively (Sicard *et al*., 2011; Wozniak *et al*., 2020). The near-isogenic lines (NILs) segregating at the (*E*)-*β*-ocimene emission QTL (QTL_1) locus were generated by introgressing QTL_1 *Cg926* allele into *Cr1504* (NILrg) and *Co1983* (NILog) through six and four rounds of backcrossing, respectively. These lines were further stabilized through 4 rounds of selfing while maintaining the QTL_1 heterozygous. The resulting lines were termed *NILrg* and *NILog*, respectively

### Plant growth conditions

Seeds were surface-sterilized, plated on half-strength Murashige-Skoog medium supplemented with gibberellic acid (Duchefa Biochemie; 3.3 mg L-1; GA) and placed for two days at 4°C. Germination took place at 22°C for ten days. To promote the transition to flowering seedlings were vernalized for ten days in 4°C. Plants were then grown in growth chambers under long-day conditions (16h light/8h dark) under a temperature cycle of 22°C during the day and 16°C during the night and at 70% humidity with a light level of 200 µmol m-2s-1.

### Scent collection

#### Headspace collection of volatile compounds emitted by *Cr* x *Cg* RILs and *Co* x *Cg* F3 population

For the purpose of previous studies (Sas *et al*., 2016; Jantzen *et al*., 2019), volatile compounds emitted by *Cr x Cg* RILs and *Co x Cg* F3 population have been collected using a dynamic headspace sampling method coupled with gas chromatography-mass spectrometry (GC-MS). The extraction, assigning, and calculation of most compound intensities was done, as described below, through a semi-automated approach. The quantification of (*E*)-*β-*ocimene emission in the *Cr* x *Cg* RILs was, however, conducted manually as described in (Sas *et al*., 2016) because the close proximity of additional compounds in the available chromatograms impaired the automatic quantification of this compound.

#### Headspace collection of volatile compounds emitted by NILs, Capsella accessions and transgenic Capsella plants

Volatile compounds emitted by NILs, *Capsella* accessions and transgenic *Capsella* plants in this study were analysed by the headspace solid phase micro extraction (HS-SPME) and gas chromatography coupled with electron impact ionization/quadrupole mass spectrometry (GC-EI-MS). HS-SPME coupled with GC-MS facilitate sample processing and improve throughput and was shown to highly efficiently capture VOCs in severals species, including those previously detected in Capsella flower blend (Flamini *et al*., 2002; Rohloff *et al*., 2005; Zorrilla-Fontanesi *et al*., 2012; Rambla *et al*., 2015; Sas *et al*., 2016; Vallarino *et al*., 2018).

Three inflorescences for each plant individual analyzed were cut and placed inside an 20 ml airtight glass vial following (Vallarino *et al*., 2018). The number of open flowers per each inflorescence was noted and later used for normalization. Up to 19 individuals were measured in one measurement sequence batch. A blank sample with no inflorescence was included in each sequence batch and unless stated otherwise, each individual was measured twice. Numbers of individuals analyzed are indicated in the respective figure legends. Samples were collected between 10:00 am and 1:00 pm and placed in a dark container for a period of approximately 30 minutes before proceeding with quantification. Time until measurement was kept approximatively constant for all samples. Sample collection was also randomized to minimize the influence of uncontrolled environmental variables. A sanity check revealed that differences in sample processing time did not affect the total amount of volatiles emitted (Kruskal-Wallis, *p* = 0.837).

Headspace samples were analyzed by solid phase micro extraction (SPME) and gas chromatography coupled with electron impact ionization/quadrupole mass spectrometry (GC-EI-MS) by using Agilent 6890N24 gas chromatograph (Agilent Technologies, Böblingen, Germany; http://www.agilent.com) and a StableFlex™ SPME-fiber with 65µm polydimethylsiloxane/divinylbenzene (PDMS-DVB) coating (Supelco, Bellefonte, USA). The headspace of samples for SPME was put under the following procedure: incubation at 45°C for 10 minutes, adsorption at 45°C for 5 minutes and desorption at 250°C for 1 minute onto a DB-624 capillary column with 60m x 0.25mm x 1.40μm film thickness (Agilent Technologies, Böblingen, Germany). The GC temperature was programmed as follows: 40°C isothermal for 2 minutes, then 10°C/min ramping to 260°C final temperature and which was then held at constant level for 10 min. The Agilent 5975B VL GC-MSD system was run using a constant flow of helium at a level of 1.0 mL/min. Desorption from the SPME fiber took place at 16.6 psi with an initial 0.1 min pulsed-pressure at 25 psi. The purge of the SPME fiber was run for 1 min at a purge flow of 12.4 mL/min (Vallarino *et al*., 2018).

### Identification and quantification of volatile compounds

The chromatograms were recorded with the mass range set to 30-300 m/z and at 20 Hz scan rate. The obtained chromatograms were controlled visually, exported in NetCDF file format using Agilent ChemStation-Software and baseline-corrected with MetAlign software (Lommen, 2009). Data processing into a standardized numerical data matrix and compound identification were done using the TagFinder software (Luedemann *et al*., 2008; Allwood *et al*., 2009). Identification of compounds was performed using mass spectral and retention time matching to the reference collection of the Golm Metabolomics Database (GMD) for volatile compounds. To assign compound as present in the analyzed sample a the criteria of at least 3 specific mass fragments per compound and a retention time deviation < 3.0% was used. Mass to charge ratios (m/z) for selective and specific quantification of annotated compounds are reported in the format Tag_mass followed by m/z (**Supplemental Data1**). To define non-annotated compounds, we correlated the mass fragments for each retention time. When mass fragments were not well correlated (r^2^ < 0.4), we performed a hierarchical clustering based on dissimilarity indices using an optimal number of clusters determined based on the silhouette method (factoextra package in R). Non-annotated compounds are named Time_###_**, with ### corresponding to the Time_group and ** to the cluster number. These yet unknown volatile compounds are characterized by assigning m/z and retention time to each Time_###_** features.

### Flower fragrance analysis

For quantification purposes, all mass features were evaluated for the best specific, selective and quantitative representation of observed analytes. The blank values of each volatile compound were subtracted from the corresponding individual samples for each sequence batch. Each sample was then standardized to the number of open flowers. For each compound at a given retention time, we selected the three best correlated (Pearson rank correlation coefficient) mass fragments and averaged blank-corrected and standardized intensities of technical replicates of each individual. Analytes present in at least two biological replicates of at least one population were kept for further analysis. The intensity of the volatile compound detected for each individual was then summed to estimate the total emission. The relative proportion of each detected compound present in the flower scent was determined by dividing each volatile compound’s intensity by the sum of the intensity of all detected compounds in the corresponding samples to obtain relative abundances of each annotated and non-annotated volatile. The comparison of *β-*ocimene emission between genotypes was performed by scaling the standardized response of the corresponding mass fragments to the maximal intensity detected in the experiment.

To assess the difference in volatile composition between *Capsella* species, we first performed a multivariate analysis on the relative proportion of each compound. We calculated Bray-Curtis dissimilarities that we illustrated with a non-metric multidimensional scaling (NMDS) plot (vegan package in R). We next performed hierarchical clustering based on the average Bray-Curtis dissimilarity scores to visualize the relationship between the volatiles profiles emitted by the different populations analyzed (Dendextend in R). We formally tested whether the mating systems affected the floral scent composition by conducting a permutational multivariate analysis of variance (PERMANOVA, adonis function from the vegan package in R using 1000 permutations) using the mating system, species and populations as factors. Additionally, we investigated the importance of different compounds for classifying populations into different mating types using the Random Forest algorithm (randomForest package in R) (Ranganathan and Borges, 2010). We use the algorithm to calculate the probability of being assigned to the correct mating systems categories based on the relative abundance of different scent compounds (n_*tree*_ = 1000, m_*try*_ = 7). We next extracted the mean decrease accuracy for each compound which describes the extent to which they contribute to the accurate classification of the samples.

Because the above analysis suggested a convergent evolution of flower scent in the two selfers, we further characterize differences in flower fragrance composition between the different species and mating system. To this end, we constructed linear models with zero-inflated beta distribution (glmmadmb package in R) using mating systems and/or species as fixed factors and population nested within mating systems or species as random variables. The model terms’ significance was determined using the likelihood test (lrtest function from the lmtest package in R). In parallel, we also conducted pairwise comparisons of the abundance of the different compounds between *Capsella* species using Tukey’s honest significant differences test to identify significantly reduced or increased compounds in the selfers compared to *C. grandiflora* (agricolae package in R). The resulting number of compounds emitted at significantly different levels by the *Capsella* species were used to visualize the overlap between the independent evolution of flower scent (VennDiagram package in R).

### QTL mapping

The genotypes of *Cr x Cg* RILs and *Co x Cg* F2 mapping populations have been obtained as previously described (Sicard *et al*., 2011; Wozniak *et al*., 2020). Scent compounds emitted by both populations were collected as described above. QTLs for the selected volatile compounds were mapped using R/QTL (Broman and Sen, 2009). The single-QTL genome scan was conducted using a non-parametric model. QTLs were tested in 1 cM intervals and assessed using LOD scores (the log10 likelihood ratio). Genome-wide permutations (1000) under the non-parametric model were used to determine significance thresholds (5%). A 1.5-LOD support interval was used to estimate the position of each QTL. The results of the QTL analyses are presented in **Table S5**. Note that our QTL analyses failed to detect significant QTL for benzeneacetaldehyde in *Co*. This could be due to a complex genetic basis involving a large number of weak effect loci also influencing these traits quantitatively and/or an important non-genetic variance limiting the statistical power to detect QTL introduced by our phenotyping approach. Indeed, here we used inflorescences of pooled F3 progenies of each F2 individual to quantify volatile abundance. In such a setup, any distortion of allele segregation in the F3 individuals could lead to an inaccurate representation of the F2 genotypes and, therefore, a considerable loss in our QTL detection power.

### Fine mapping

We first identified a *NILrg* segregating in the region containing the QTL_1 interval by genotyping plants with markers G06, G07 and G09 (**Table S7**). To refine the position of QTL_1, around 1,200 *NILrg* progenies were screened for plants harbouring a meiotic recombination between the markers m1621 and m1629, each located at the opposed borders of the QTL confidence interval (**Table S5**). The position of the recombination breakpoints was determined by genotyping informative recombinants with additional markers located in the focal region (**Table S7**). Floral scent was collected from each of the selected recombinant individuals two times using the static headspace sampling method as described above.

### Molecular cloning and plant transformation

*Cr* (*TPS02r*) and *Cg* (*TPS02g*) genomic constructs were used to complement the *NIL* plants homozygous for the *Cr* allele (*NILrr*). Both constructs were PCR-amplified from either *NILrr* or *NILgg* (NIL homozygous for the *Cg* allele) genomic DNA. A 6.1 kb *TPS02r* and 5.9 kb *TPS02g* genomic fragments starting ∼1.3 kb before the transcription start site of *TPS02* and ending ∼1.3kb after the stop codon were amplified in two PCR reactions as described in **Table S8** (sequences of primers used are presented in **Table S9**). The genomic fragments were assembled and subcloned into a modified version of pBluescript II KS (StrataGen, pBlueMLAPUCAP) using the InFusion® HD Cloning Plus kit (Clontech), yielding the construct pBS-TPS02r and pBS-TPS02g. Following the same cloning approach, a sequence coding for the yellow fluorescent protein was inserted before the codon stop position of TPS02 in the resulting plasmids. The sequences corresponding to the TPS02YFP recombinant proteins were then transferred to pAS95, modified pBluescript II KS containing a CaMV 35S promoter and terminator, yielding the 35S::TPS02gYFP and the 35S::TPS02rYFP constructs. The serine at the 8^th^ amino acid position was then converted into a proline by PCR-based site-directed mutagenesis, giving rise to the 35S::TPS02g^S8P^YFP construct. Details about the fragment used to generate the different constructs are given in **Table S8**.

After verification by Sanger sequencing, the fragments were transferred into the AscI site of the plant transformation vector pBarMAP, a derivative of pGPTVBAR (Becker *et al*., 1992). These genomic constructs were then used to transform *Capsella NILrr* plants by floral dip (Clough and Bent, 1998) or Arabidopsis thaliana leaf protoplasts using PEG-mediated transfection (Wu *et al*., 2009).

### Gene expression and splicing analysis

Total RNA was extracted from inflorescences using Trizol (Life Technologies). RNAs were treated with TurboDNAse (Ambion) and reverse transcribed using oligo(dT) and the Superscript III Reverse Transcriptase (Invitrogen). Quantitative RT-PCR was performed using the SensiMix SYBR Low-ROX kit (Bioline) and a LightCycler® 480 (Roche). The primers used are described in **Table S9**. For each genotype, three biological replicates were used, and for each of them, three technical replicates were included. To detect splicing difference, primers onw540 and onw541 were used to amplify the full-length *TPS02* transcripts.

### Confocal imaging

To analyze the cellular localization of TPS, we imaged transformed *A. thaliana* protoplast with a confocal laser scanning microscope Zeiss LSM710 (Zeiss) using an excitation wavelength of 514 nm, with emission collected between above 630 nm for Chlorophyll A and between 520 - 570 for YFP. Above-threshold Pearson’s correlation coefficient between the Chloroplhyll A and YFP signal intensities were determined using ImageJ (http://rsbweb.nih.gov/ij/) to assess the efficiency of TPS02 targeting within chloroplasts.

### Sequence and population-genetic analyses

Pairwise sequence alignments of nucleotide sequences of *NILrr* and *NILgg* TPS02 alleles was performed using EMBOSS Needle (Rice *et al*., 2000). Protein sequence alignments were performed using the Clustal Omega and visualized by MView (Brown *et al*., 1998; Rice *et al*., 2000). Chloroplast transit peptide sequence was predicted using the LOCALIZER 1.0.4 (Sperschneider *et al*., 2017).

To correlate the genotype at candidate polymorphisms with scent emission, five *Cg* individuals and five *Cr* accessions (three individuals per each accession) were sequenced in the regions containing five non-synonymous variants within *TPS02* and phenotyped for (*E*)-*β-*ocimene emission as mentioned above.

The dataset used for the population genetic analysis included 180 Cg individuals from a single population and species-wide samples of 13 *Cg* and, 51 *Cr* and 16 *Co* individuals (Ågren *et al*., 2014; Josephs *et al*., 2015; Koenig *et al*., 2018). TPS02 haplotypes were reconstituted by a combination of local assembly and multiple pair-end phasing (similarly to (Sicard *et al*., 2016)). The resulting sequences were used to reconstruct neighbour-joining trees with MEGA11 using the Tamura 3-parameter method and gamma distribution (Tamura *et al*., 2021). MEGA11 was also used to compute genetic distances as the number of nucleotide differences per sequence comparison. The hierarchical clustering of the samples based on the genotypes at *Cr1504*/*Cg926* TPS02 non-synonymous variants was done using Elucidean distances. The count of *Cr1504* or *Crub183* k-mers of different sizes in different samples were estimated using JELLYFISH (Marçais and Kingsford, 2011).

To visualise the genetic distances between the three *Capsella* species, reads were mapped to the *Capsella* reference genome (Slotte *et al*., 2013) using Stampy v1,0,19 (Lunter and Goodson, 2011). After bioinformatics processing using Picard tools (https://picard.sourceforge.net/), Raw SNP calls were generated by the joint calling of all individuals in GATKv4 (Van der Auwera and O’Connor, 2020). To determine the genetic relationship between *Capsella* species, we first identify the ten most unrelated individuals from each species. For each species separately, we pruned the dataset with a target of ten individuals by varying the relatedness cutoff using plink 1.9 (Chang *et al*., 2015). We then computed the distance matrix with the same software using the “hamming distance” procedure. The dendrogram was created by performing a hierarchical clustering based on the resulting distance matrices using the Ward method in the R package *hclust*.

*C. rubella* phylogeny was reconstituted using the substitution model in the IQTREE software (Nguyen *et al*., 2015). To root the tree, we used a *C. grandiflora* individual representative of the species’ genetic identity, i.e. the individual with the lowest contribution on the principal components of a genetic principal component analysis performed with the R package *pcadapt* (Luu *et al*., 2017). We looked at the *TPS02* haplotype diversity across *C. rubella* genotypes by extracting the variants located in the *TPS02* locus and performing an admixture analysis (Alexander *et al*., 2009). To further assess the regime of selection under which TPS02 evolves, we performed a whole genome screen of synonymous versus non-synonymous polymorphism (π_S_ and π_N_ respectively) in coding regions. Based on the reference genome sequences, we defined the 4-fold degenerate nucleotide positions as synonymous sites (S sites) and the 0-fold degenerate nucleotide positions as non-synonymous sites (N sites). We compute the nucleotide diversity in S and N sites per gene and per species using customized r script and the *nuc*.*div* function from *pegas* R package (Paradis, 2010).

### Statistical analysis

Statistical analyses were conducted in R statistical programming language (R Core Team, 2018). For multiple comparisons of phenotypic means, Tukey-HSD tests were performed using agricolae package add-ons implemented in R for multi-comparison tests. Data were presented as mean and standard deviation and comparisons with p values below 0.05 were considered significant. Data were illustrated as box plots using the ggplot2 package implemented in R (R Core Team, 2018). Outliers were defined as values deviating from the mean by more than 3 Standard deviations. The lower and upper hinges in all boxplots correspond to the first and third quartiles, respectively. The upper and lower whiskers add or subtract 1.5 interquartile ranges to/from the 75 and 25 percentile, respectively. The middle lines represent the median.

## Acknowledgments

We thank Tanja Slotte and Barbara Neuffer for seeds; Fredric Hedlund and Kathrin Hesse for plant care, and members of the Lenhard, and Rosa groups for discussion and comments on the article. The computation and data handling were provided by the Swedish National Infrastructure for Computing (SNIC) at Uppmax, partially funded by the Swedish Research Council through grant agreement no 2018-05973. This work was supported by the Deutsche Forschungsgemeinschaft (grant SI1967/2) and the Swedish Research Council (grant 2018-04214) to AS and a Carl Tryggers Stiftelse (CTS 18-350) and Marie Skłodowska-Curie Individual Fellowships (MSCA-IF 101032710) to LZ.

## Author contributions

NW, KS, JK, ML and AS designed research. NW, LZ, IF and FJ, performed experiments. KS, MO and CK performed the population genetic analyses, and AE carried out the metabolic profiling. AS and JK supervised the project. NW, KS and AS wrote the manuscript with help from all authors. All authors discussed and commented on the manuscript.

## Figure legends

**Figure S1:**
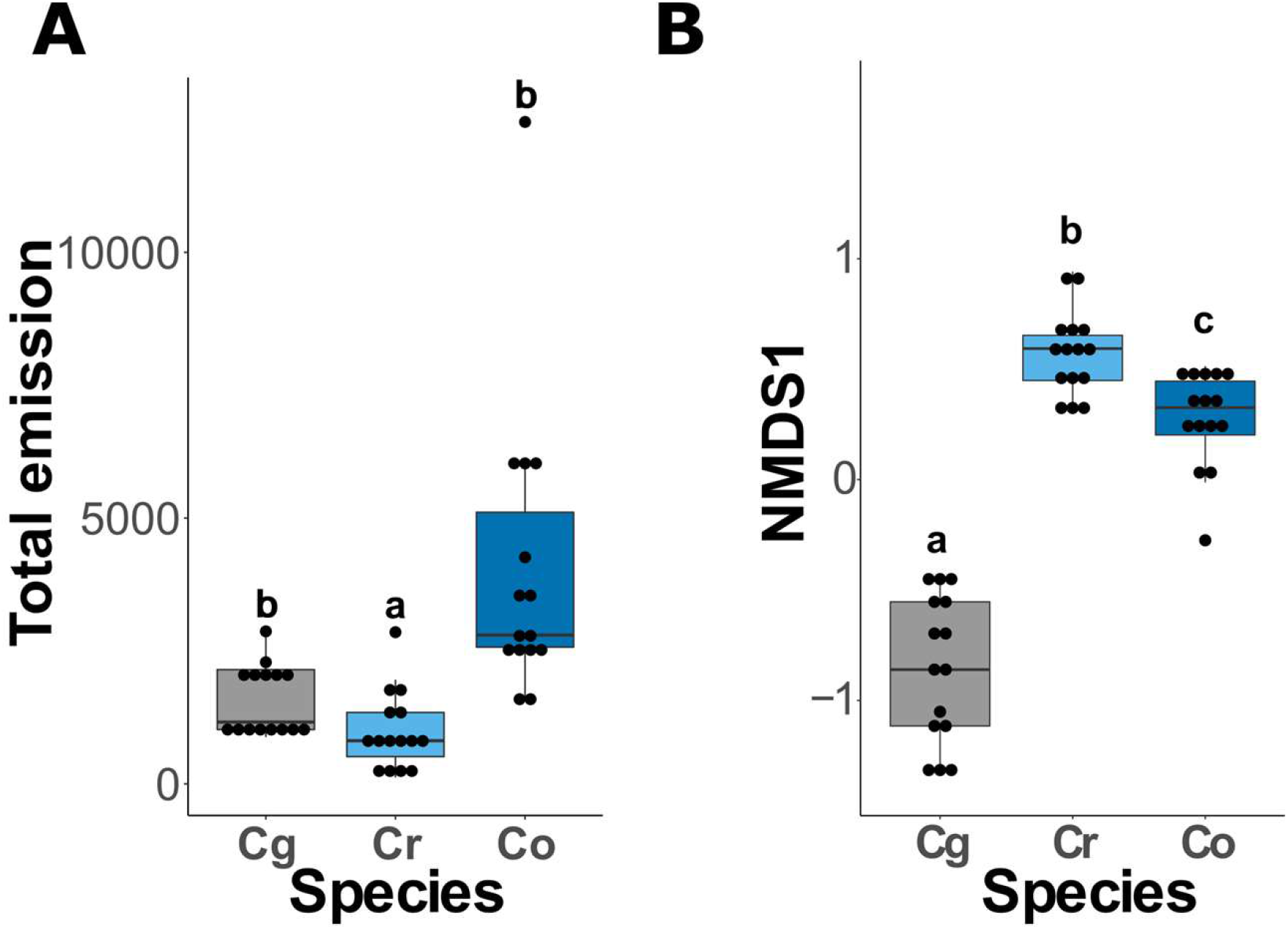
Convergent evolution in the composition of flower scent between independent transitions to selfing in the genus *Capsella*. (**A**) Total volatile emission detected in the different *Capsella* species (n =15 per species) (**B**) Distribution of NMDS1 scores among *Capsella* species (n = 15 per species). Letters illustrate significant differences as determined by one-way ANOVA with Tukey’s HSD test (α=0.05).

**Figure S2:**
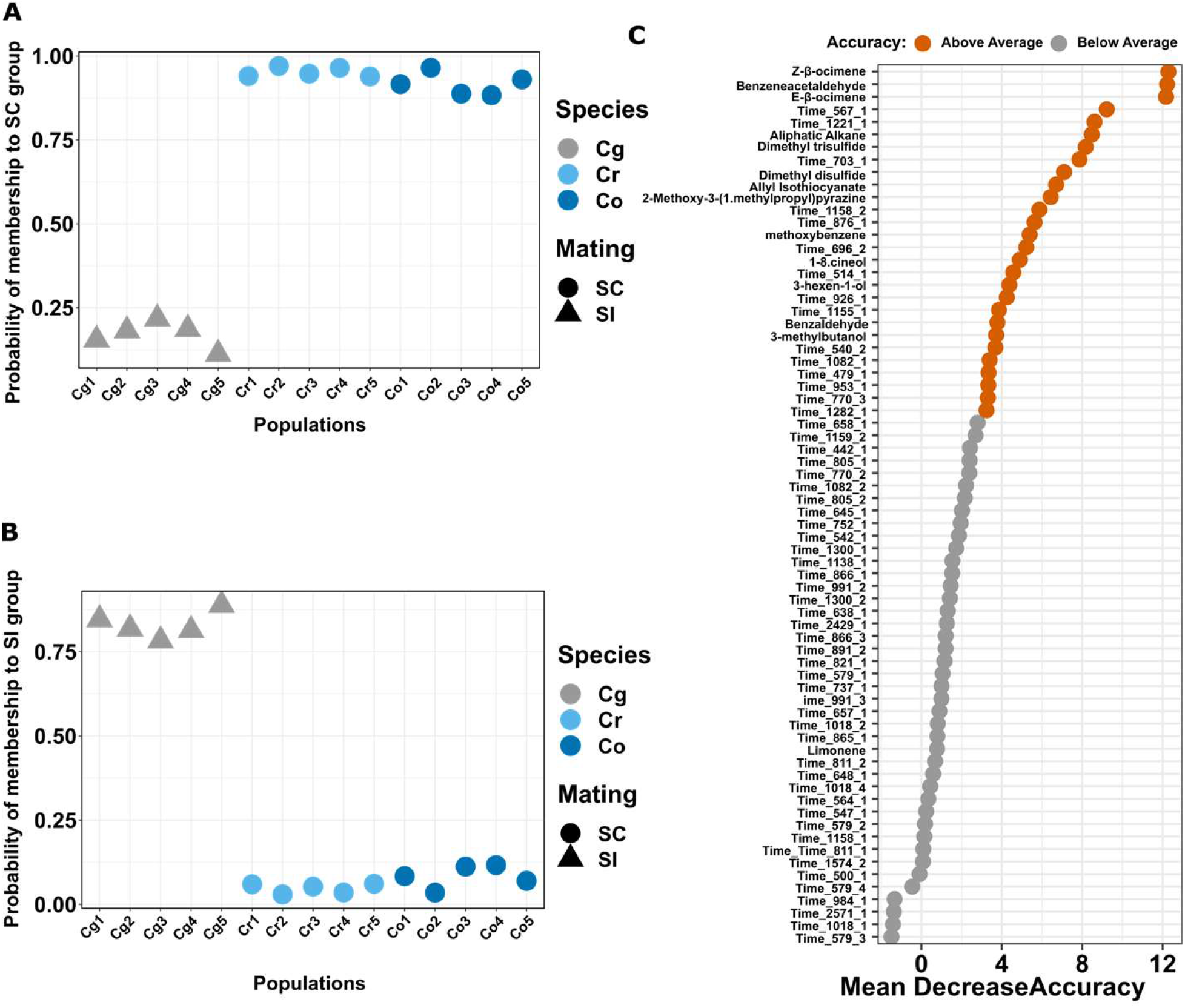
The composition of flower fragrance predicts mating system. (**A**) The “out of the bag” probability of the different populations to be classified as self-compatible by the random forest model. (**B**) The “out of the bag” probability of the different populations to be classified as self-incompatible by the random forest model. (**C**) Random forest variable importance shown as the Mean Accuracy Decrease in model prediction caused by dropping each variable

**Figure S3:**
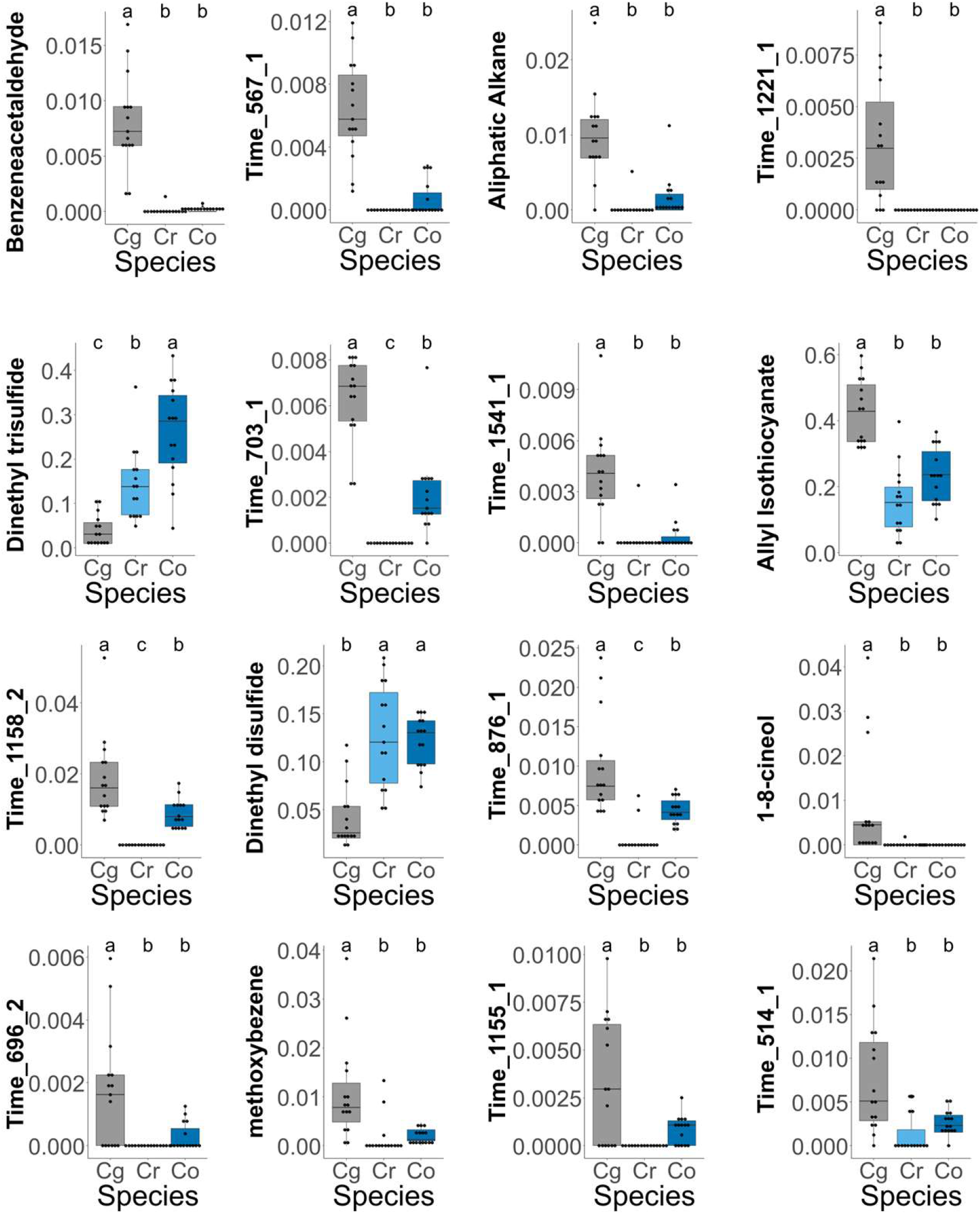
Similarity in the emission of volatile compounds between independently evolved selfing lineages. The relative abundance of the compounds in each *Capsella* species (n=15 for each species) found to be significantly affected by the mating system is shown. Letters indicate significant differences as determined by one-way ANOVA with Tukey’s HSD test) (α=0.05).

**Figure S4:**
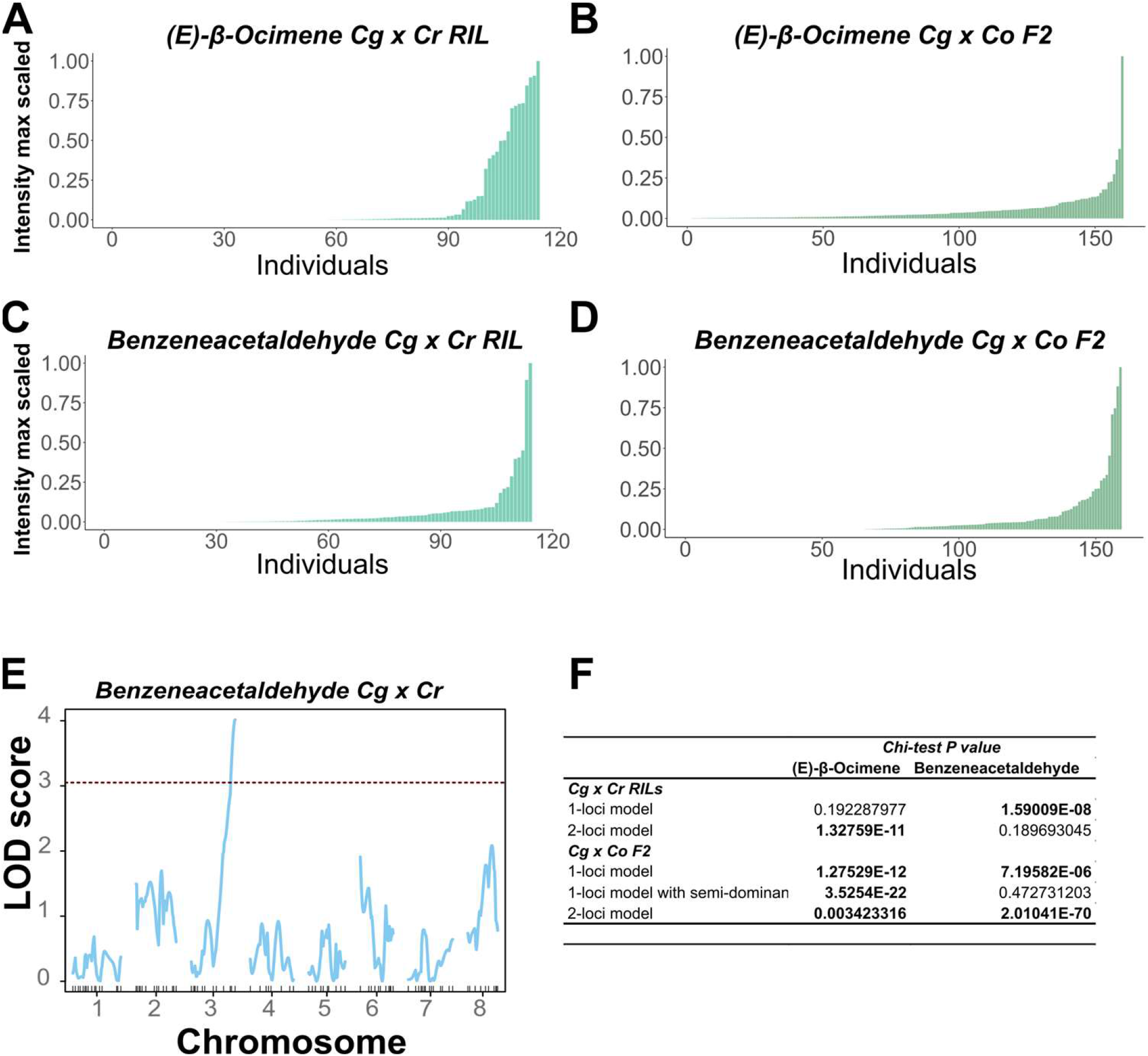
Genetic basis of the loss of volatile emission in the *Capsella* selfer species. (**A**) to (**D**) (E)-β-ocimene (**A** and **B**) or benzeneacetaldehyde (**C** and **D**) emission intensity scaled to maximum emission in the experiment in the *Cr1504* x *Cg926* RIL (**A** and **C**) or *Co1983* x *Cg926* F2 (**B** and **D**) populations. (**E**) QTL mapping for the emission of benzeneacetaldehyde (Tag_mass 120) in the *Cr1504* x *Cg926* RIL population. Red dashed lines indicate the genome-wide 5% significance threshold. (**F**) Comparison between observed and expected segregation ratios.

**Figure S5:**
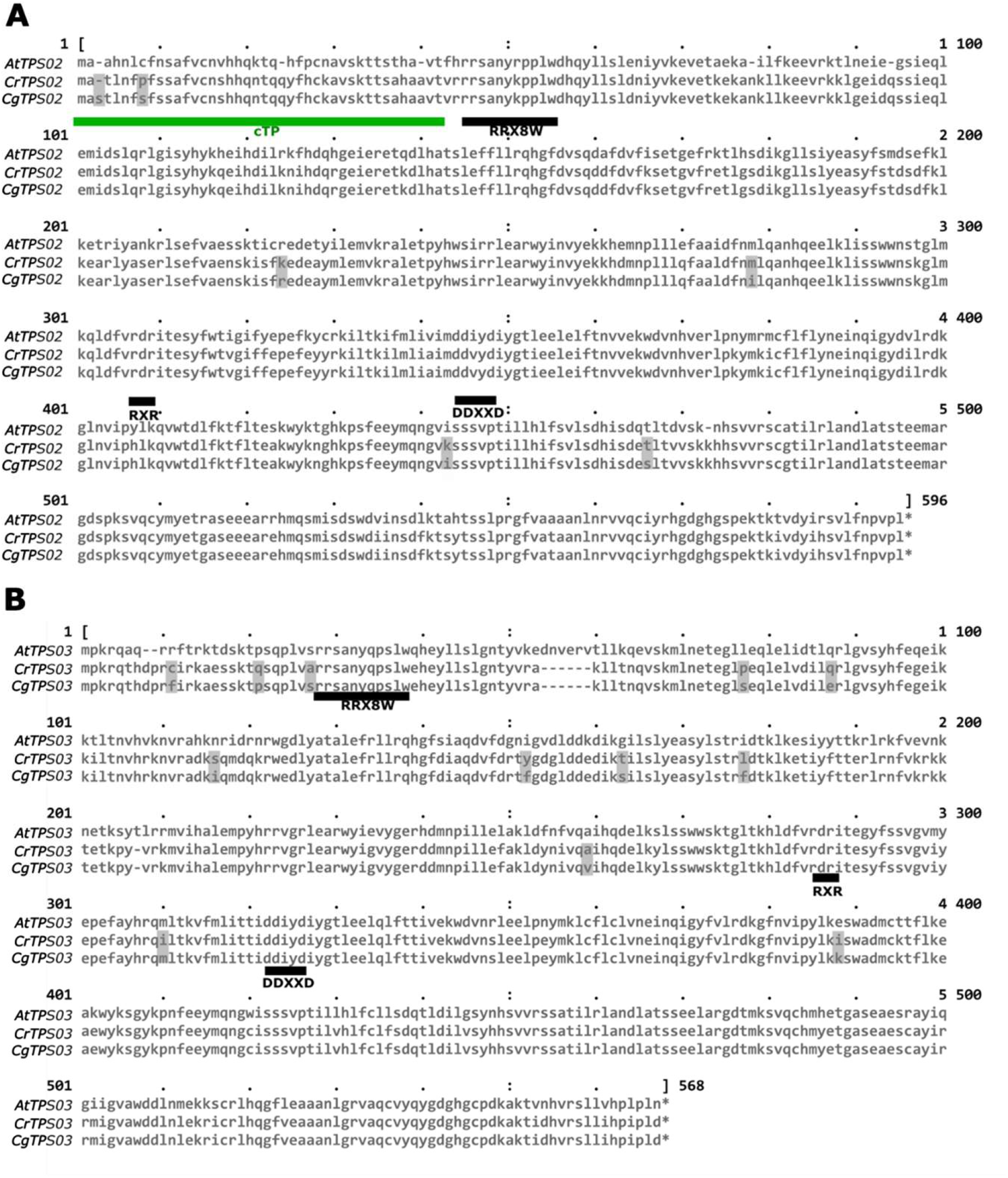
Comparison of TPS02 and TPS03 protein sequences in *C. grandiflora* and *C. rubella*. **(A)** Amino acid sequence alignment of the full-length *A. thaliana, C. rubella* and *C. grandiflora* TPS02 and **(B)** TPS03 proteins. Amino acid differences between *C. rubella* and *C. grandiflora* proteins are shaded in gray. Dashes indicate gaps. Horizontal black lines indicate the highly conserved RRX8W, RXR and DDXXDD motifs. Horizontal green line indicates the chloroplast transit peptide sequence.

**Figure S6:**
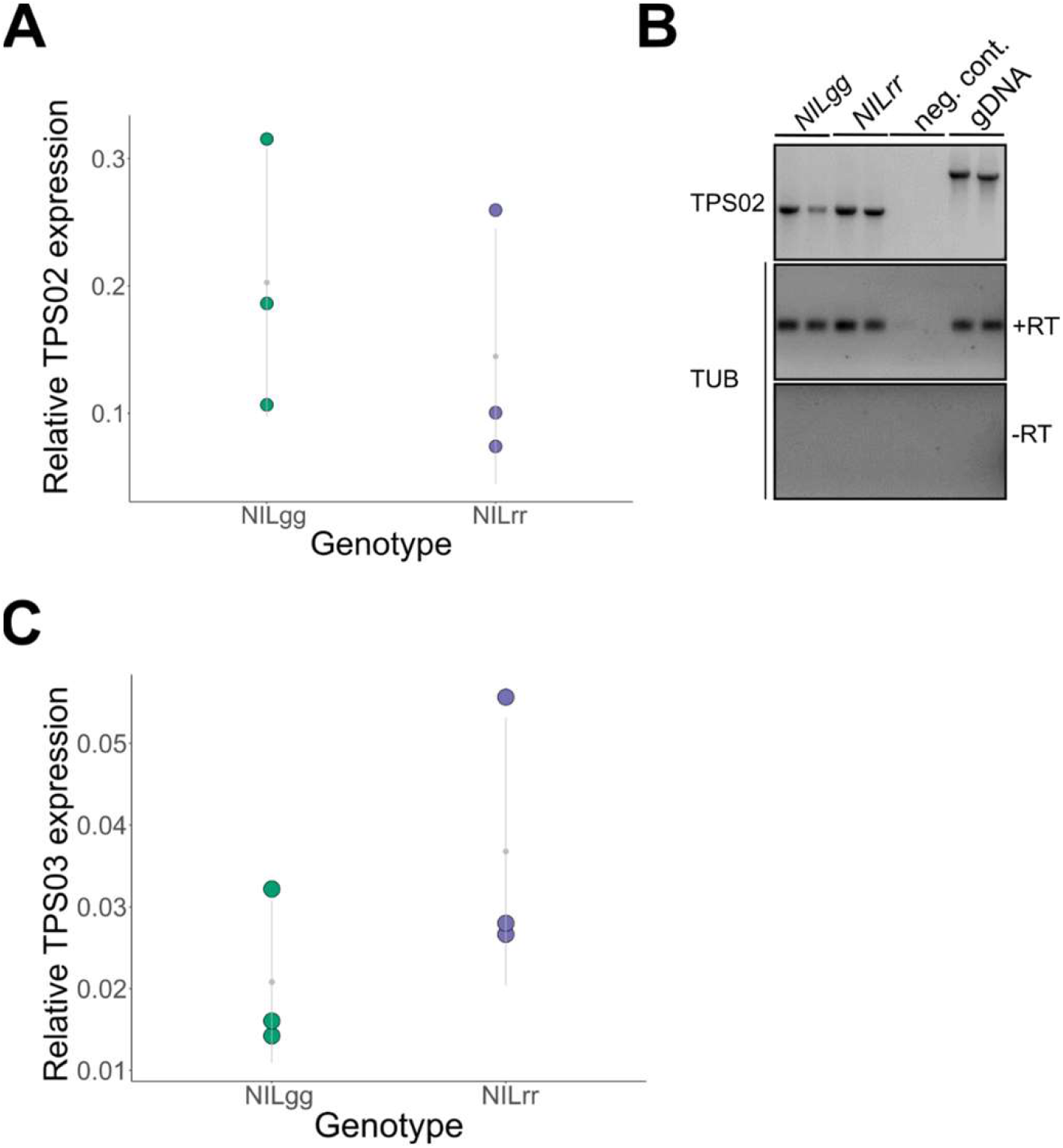
Expression of TPS02 and TPS03 is not affected in *C*.*rubella*. (**A**) and (**C**) Relative expression of *TPS02* (**A**) and *TPS03* (**C**) in NIL homozygous for the *Cg* (NILgg) or *Cr* (NILrr) *TPS02* allele determined by quantitative reverse transcription–PCR normalized to *Capsella TUB6*. Each dot represent the average value of each biological replicate. (B) Electrophoresis of PCR products of the full length *TPS02* ORFs or *Capsella TUB6* illustrating the fact that both alleles expressed *TPS02* mRNA of identical size. NILsrr and NILgg cDNA or genomic DNA (gDNA) were used as template. Negative PCR (neg. cont.) and reverse transcription (-RT) controls are also shown. Note that the –RT control panel does not contain positive gDNA control but was performed in parallel as the above panel.

**Figure S7:**
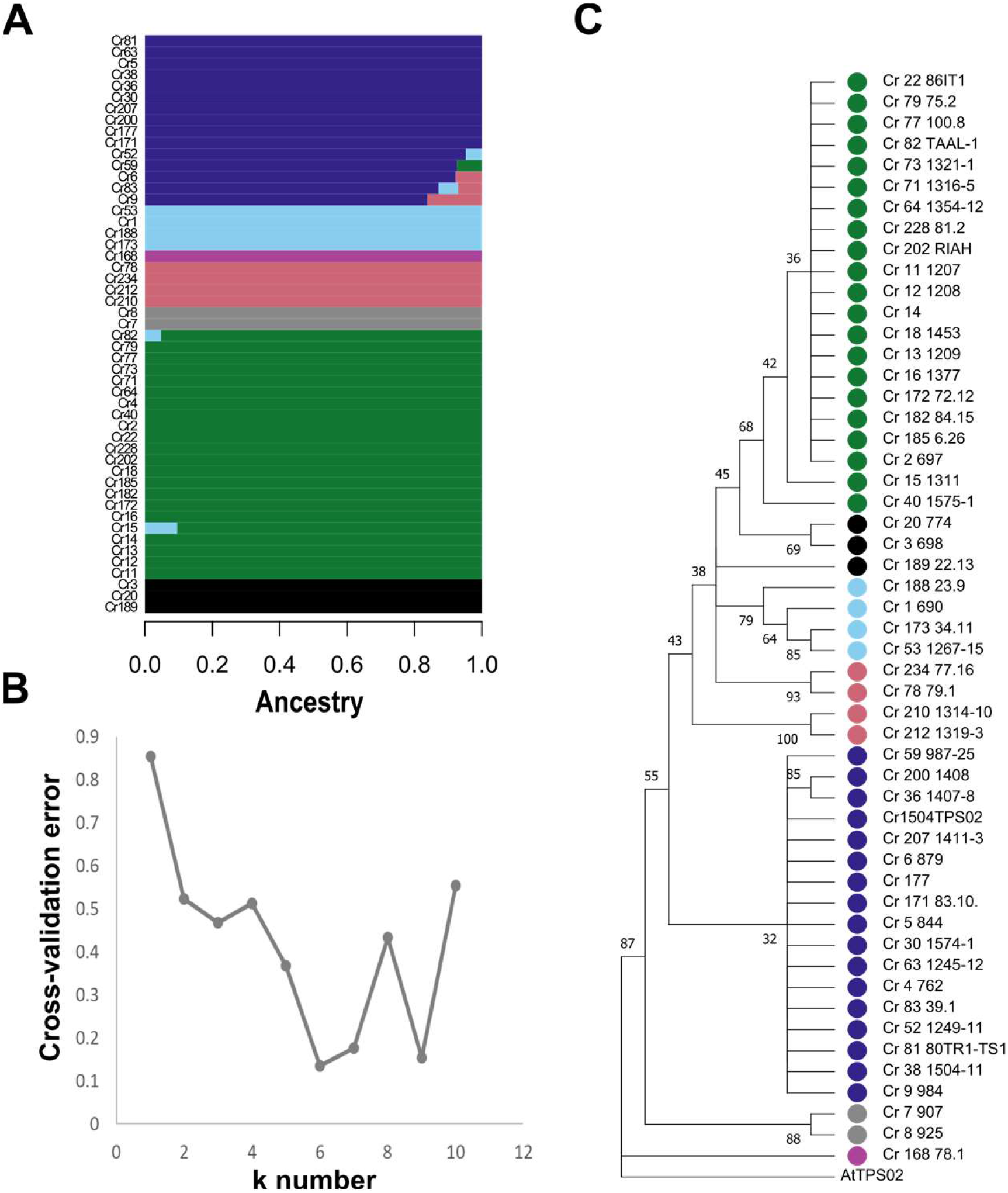
Population genetic structure and phylogenetic relationship at TPS02 among 51 *C. rubella* accessions. (**A**) ADMIXTURE plot for *C. rubella TPS02*. Each accession is represented by an horizontal line partitioned into colored segments. The size of the segment illustrates the proportion of membership to 6 clusters. (**B**) Cross validation error calculated for different admixture model (k = 0 to 10). (**C**) Evolutionary history of Cr TPS02 inferred using the UPGMA clustering. The bootstrap consensus tree was inferred from 1000 replicates. Branches corresponding to partitions reproduced in less than 60% bootstrap replicates are collapsed. The percentage of replicate trees in which the investigated accession clustered together are shown next to the branches. The color symbols at the end of each branch represent the membership to the different k clusters.

**Figure S8:**
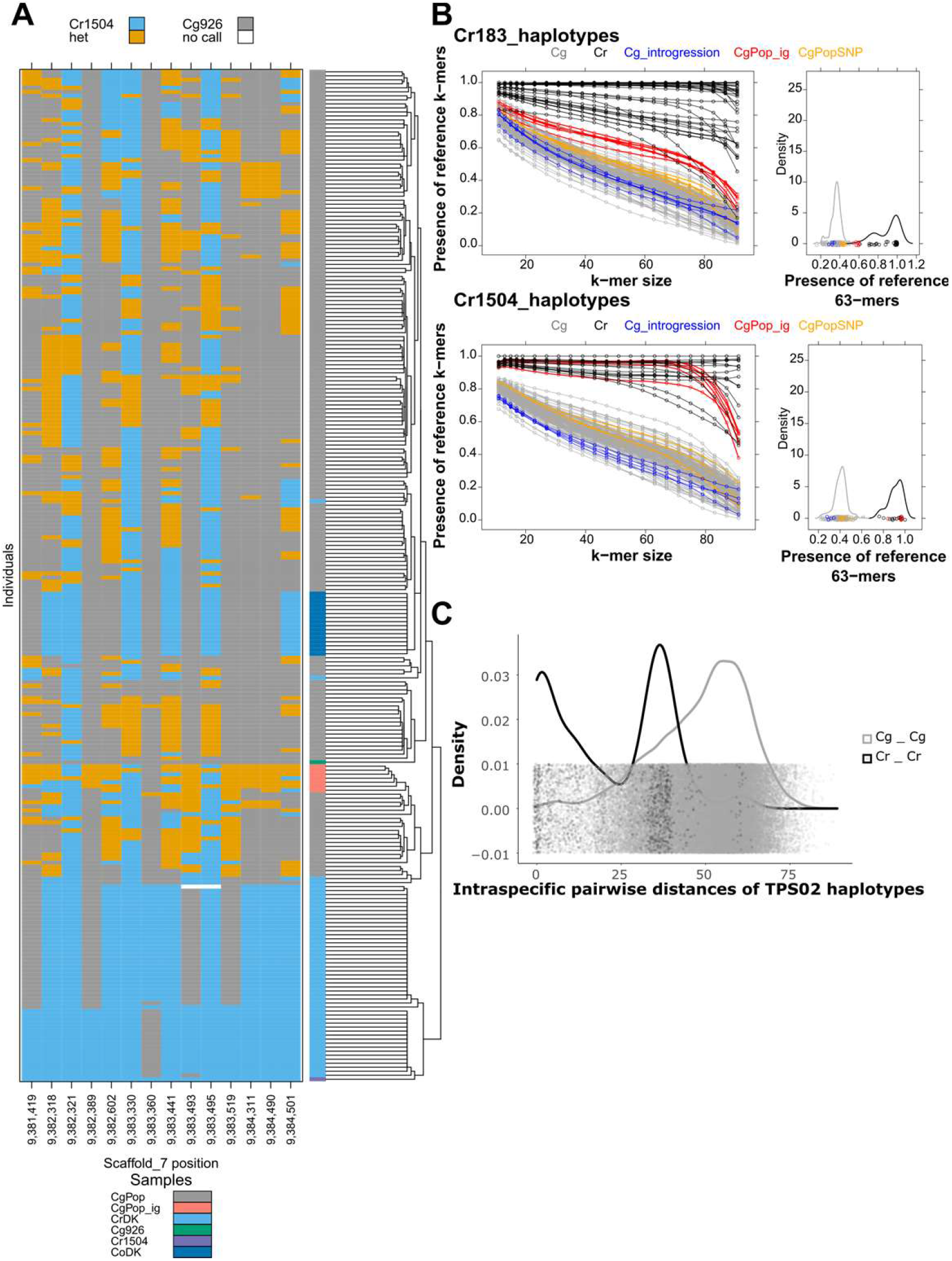
Population Genetic structure and phylogenetic relationship at TPS02 among *Capsella* species. (**A**) Haplotype clustering and genotype at the candidate polymorphisms in *TPS02* of all the resequenced *Capsella* individuals, including all publicly available *Cg* (CgPop), *Cr* (CrDK) and *Co* (CoDK) genomes as well as of the parental *TPS02g* (*Cg926*) and *TPS02r* (*Cr1504*) alleles. *Cg* samples showing sign of recent intogression at TPS02 locus (CgPop_ig) are shown in red. (**B**) Introgression events were estimated by counting the presence of *Crub183* (upper panel) or *Cr1504* (lower panel) like k-mers of increasing length in genome re-sequencing raw data. Each line represents an individual. Individuals with putative introgression were identified based on the distribution of the presence of 63-mers in *Cr* or *Cg* shown on the right panels. All individuals with outlier values (apart from the mean by more than 3 SD) in the distribution of their respective species were considered to carry an introgressed allele at TPS02. *Cr* individuals with introgression from *Cg* (Cg_introgression) are shown in blue, *Cg* individuals with introgression from *Cr* (CgPop_ig) are shown in red, *Cg* individuals harboring the *Cr* allele at the candidate causal at SNP 9.384.490 are displayed in orange the remaining *Cg* and *Cr* individuals are indicated in gray and black, respectively. (**C**) Density distribution of pairwise distances in number of polymorphisms between *Cr* (Cr-Cr) or *Cg* (Cg-Cg) *TPS02* haplotypes.

**Table S1:**
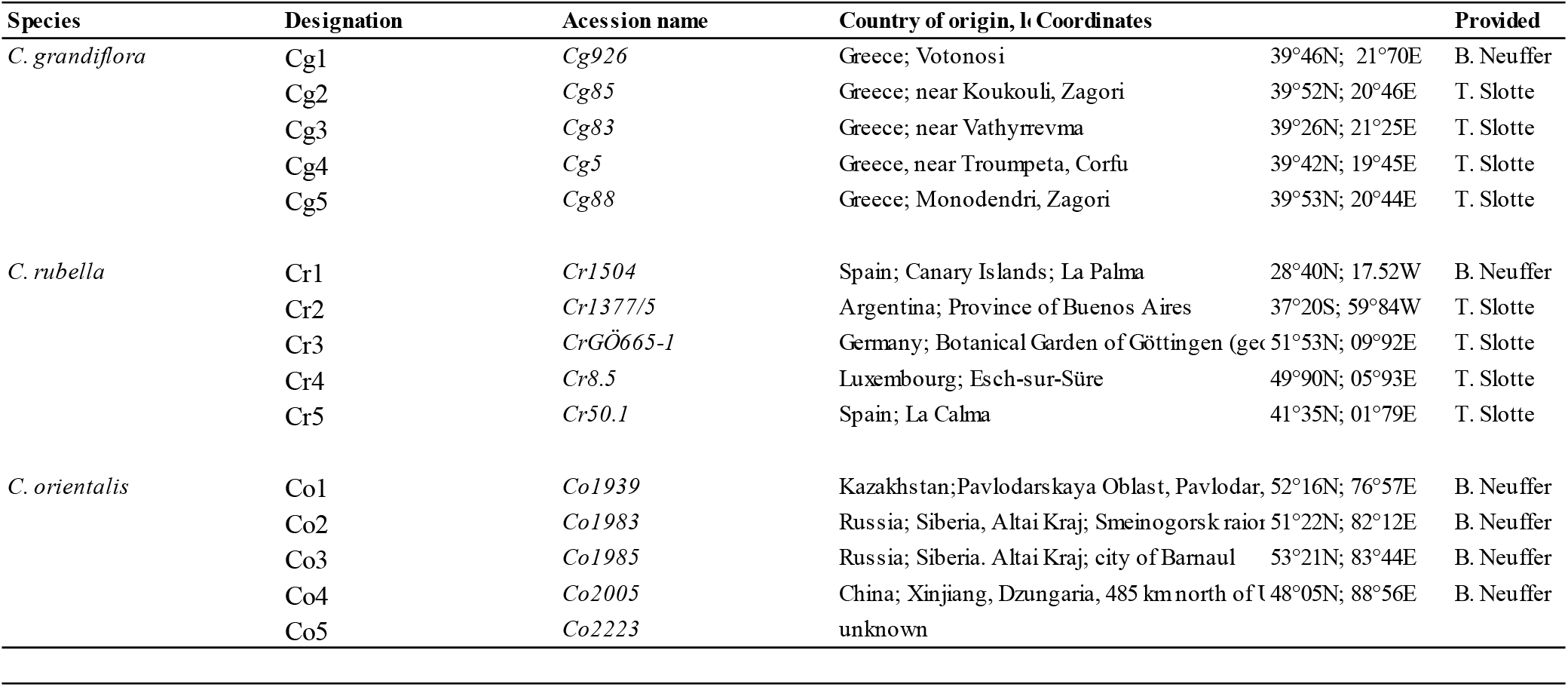
Geographical origin of the Capsella populations used in this study.

**TableS2:**
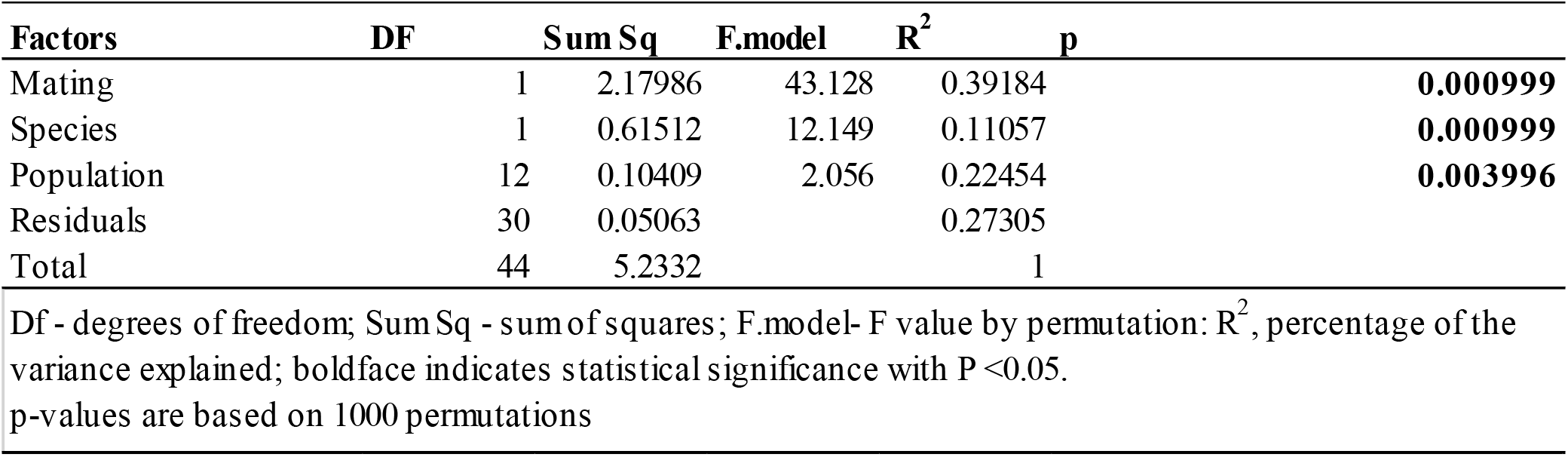
PerMANOVA results based on Bray-Curtis dissimilarities using volatile compounds proportion.

**Table S3:**
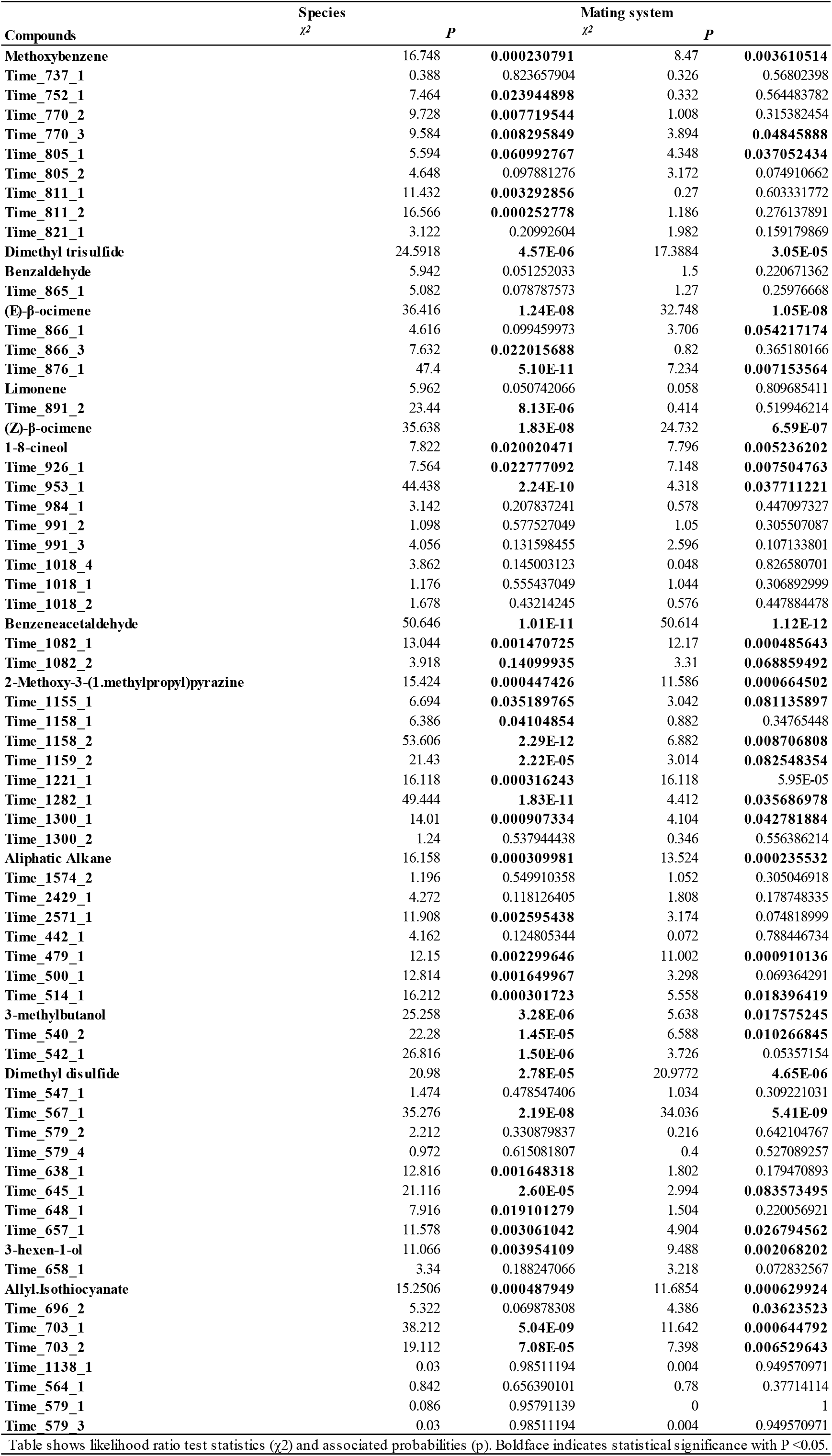
Summary of Likelihood Ratio Test comparing nested generalized linear models assessing the fixed effect of Species and.

**TableS4:** Summary statistics of volatile emission in the different Capsella species.

| Compounds | Cg |  | Species<br>Cr |  | Co |  |
| --- | --- | --- | --- | --- | --- | --- |
|  | M | S | M | S | M | S |
| Methoxybenzene | **0.010630783** | **a** | **0.001677217** | **b** | **0.001982741** | **b** |
| Time_737_1 | 0.018515338 | a | 0.028106195 | a | 0.016252878 | a |
| Time_752_1 | 0.003590433 | ab | 0.000391577 | b | 0.008283584 | a |
| Time_770_2 | 0.001603701 | b | 0.013827992 | a | 0.002323329 | b |
| Time_770_3 | 0.00160566 | a | 0.000972872 | a | 0.000862741 | a |
| Time_805_1 | 0.00352704 | b | 0.031024329 | a | 0.00952335 | b |
| Time_805_2 | 0.012344405 | b | 0.100079685 | a | 0.036189582 | b |
| Time_811_1 | 0.00119366 | a | 0.000590627 | a | 0.002143058 | a |
| Time_811_2 | 0.001189849 | a | 0.000734592 | a | 0.002544222 | a |
| Time_821_1 | 0.002835805 | a | 0.000517721 | b | 0.001348099 | ab |
| Dimethyl trisulfide | **0.038688425** | **c** | **0.140618335** | **b** | **0.258357857** | **a** |
| Benzaldehyde | 0.002397205 | a | 2.22E-10 | b | 0.000870513 | ab |
| Time_865_1 | 0.002769065 | a | 0.00228266 | a | 0.001870321 | a |
| (E)-β-ocimene | **0.030639656** | **a** | **9.16E-05** | **b** | **0.000820914** | **b** |
| Time_866_1 | 0.009997283 | a | 0.004404194 | ab | 0.001339148 | b |
| Time_866_3 | 0.001505189 | a | 0.005627841 | a | 0.003462713 | a |
| Time_876_1 | **0.00964332** | **a** | **0.000695227** | **c** | **0.004355748** | **b** |
| Limonene | 0.007726969 | ab | 0.005079187 | b | 0.01289595 | a |
| Time_891_2 | 0.001959857 | ab | 0.000475104 | b | 0.004869335 | a |
| (Z)-β-ocimene | **0.065674771** | **a** | **2.22E-10** | **b** | **0.001922487** | **b** |
| 1-8-cineol | **0.008159534** | **a** | **0.000122407** | **b** | **2.22E-10** | **b** |
| Time_926_1 | 0.007881806 | a | 0.009449743 | a | 0.001196539 | a |
| Time_953_1 | 2.22E-10 | b | 2.22E-10 | b | 0.014350838 | a |
| Time_984_1 | 0.002055672 | a | 0.000901936 | a | 0.001741308 | a |
| Time_991_2 | 0.001098289 | b | 0.006469845 | a | 0.000819688 | b |
| Time_991_3 | 0.00179305 | b | 0.011217214 | a | 0.002402198 | b |
| Time_1018_4 | 0.000999395 | a | 0.002252073 | a | 0.00208052 | a |
| Time_1018_1 | 0.005075483 | a | 0.011984793 | a | 0.006370617 | a |
| Time_1018_2 | 0.000788519 | b | 0.003404677 | a | 0.001013737 | ab |
| Benzeneacetaldehyde | **0.007963764** | **a** | **8.81E-05** | **b** | **7.87E-05** | **b** |
| Time_1082_1 | 0.001159067 | b | 0.008469001 | a | 0.006229031 | ab |
| Time_1082_2 | 0.000198336 | b | 0.004720871 | a | 0.002189575 | ab |
| Time_1138_1 | 0.000252564 | a | 0.001359183 | a | 0.001756782 | a |
| 2-Methoxy-3-(1-methylpropyl)pyrazine | 0.002193157 | b | 0.014706115 | a | 0.005738541 | b |
| Time_1155_1 | **0.003240209** | **a** | **2.22E-10** | **b** | **0.000840797** | **b** |
| Time_1158_1 | 0.000884318 | a | 0.001808006 | a | 0.002862705 | a |
| Time_1158_2 | **0.018428539** | **a** | **2.22E-10** | **c** | **0.008651478** | **b** |
| Time_1159_2 | 8.82E-05 | b | 2.22E-10 | b | 0.002598038 | a |
| Time_1221_1 | **0.003164588** | **a** | **2.22E-10** | **b** | **2.22E-10** | **b** |
| Time_1282_1 | 2.22E-10 | b | 2.22E-10 | b | 0.005168853 | a |
| Time_1300_1 | 0.000373914 | a | 0.001955918 | a | 0.001408788 | a |
| Time_1300_2 | 0.00060735 | a | 0.00226837 | a | 0.001054737 | a |
| Aliphatic Alkane | **0.009788108** | **a** | **0.000334788** | **b** | **0.00155746** | **b** |
| Time_1574_2 | 0.004392946 | a | 0.004685916 | a | 0.001812817 | a |
| Time_2429_1 | 0.001373451 | b | 0.011450187 | a | 0.000620092 | b |
| Time_2571_1 | 0.002443452 | ab | 0.006689068 | a | 0.000771463 | b |
| Time_442_1 | 0.000968872 | b | 0.007009022 | a | 0.000488195 | b |
| Time_479_1 | 0.002609063 | a | 0.002312167 | a | 2.22E-10 | b |
| Time_500_1 | 0.026719228 | a | 0.005513139 | a | 0.007544806 | a |
| Time_514_1 | **0.007401431** | **a** | **0.001220439** | **b** | **0.002581625** | **b** |
| 3-methylbutanol | 0.007738506 | b | 0.037124784 | a | 0.011337656 | b |
| Time_540_2 | 0.012513894 | b | 0.03921192 | a | 0.019220535 | b |
| Time_542_1 | 0.000187018 | b | 0.000303767 | b | 0.003174817 | a |
| Dimethyl disulfide | **0.041465637** | **b** | **0.12537832** | **a** | **0.120451848** | **a** |
| Time_547_1 | 0.001980775 | b | 0.021522309 | a | 0.00292954 | b |
| Time_564_1 | 0.003044396 | a | 0.001460793 | a | 0.000849583 | a |
| Time_567_1 | **0.006242829** | **a** | **2.22E-10** | **b** | **0.000685327** | **b** |
| Time_579_1 | 0.001116466 | a | 0.001505966 | a | 0.0007846 | a |
| Time_579_2 | 0.006203741 | b | 0.015983086 | a | 0.005178891 | b |
| Time_579_4 | 0.001036258 | a | 0.002954909 | a | 0.001113768 | a |
| Time_579_3 | 0.001197008 | a | 0.001254646 | a | 0.000849304 | a |
| Time_638_1 | 0.000240806 | b | 2.22E-10 | b | 0.002285104 | a |
| Time_645_1 | 8.43E-05 | b | 2.22E-10 | b | 0.007871635 | a |
| Time_648_1 | 0.00402021 | b | 0.01458094 | a | 0.008557778 | ab |
| Time_657_1 | 0.000888302 | a | 0.002848489 | a | 0.002638806 | a |
| 3-hexen-1-ol | 0.022188929 | b | 0.071869069 | a | 0.041935606 | b |
| Time_658_1 | 0.004999163 | b | 0.018780382 | a | 0.010073768 | b |
| Allyl Isothiocyanate | **0.417271045** | **a** | **0.154120634** | **b** | **0.231652172** | **b** |
| Time_696_2 | **0.001671301** | **a** | **2.22E-10** | **b** | **0.000277054** | **b** |
| Time_703_1 | **0.006186394** | **a** | **2.22E-10** | **c** | **0.002080995** | **b** |
| Time_703_2 | **0.087360047** | **a** | **0.01128777** | **c** | **0.047680496** | **b** |

M indicates the mean proportion of different volatile compounds within the flower fragrance of different Capsella species, S show significant differences as determined by one-way ANOVA with Tukey's HSD test.

**Table S5:**
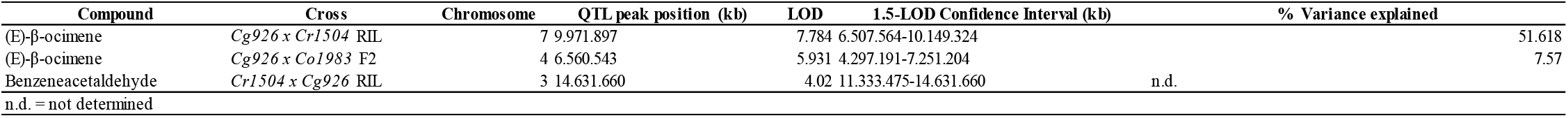
Position of the quantitative trait loci controlling the emission of the compounds the most differentiated between Capsella mating types (selected Tag_mass: 93 for (E)-β-ocimene 120 for benzeneacetaldehyde)

| Compound | Cross | Chromosome | QTL peak position (kb) | LOD | 1.5-LOD Confidence Interval (kb) | % Variance explained |
| --- | --- | --- | --- | --- | --- | --- |
| (E)-β-ocimene | *Cg926* x *Cr1504* RIL | 7 | 9.971.897 | 7.784 | 6.507.564-10.149.324 | 51.618 |
| (E)-β-ocimene | *Cg926* x *Co1983* F2 | 4 | 6.560.543 | 5.931 | 4.297.191-7.251.204 | 7.57 |
| Benzeneacetaldehyde | *Cr1504* x *Cg926* RIL | 3 | 14.631.660 | 4.02 | 11.333.475-14.631.660 | n.d. |

n.d. = not determined

**Table S6:** In silico predictions of transit peptide by localizer 1.0.4.

|  | Chloroplast | Mitochondria | Nucleus |
| --- | --- | --- | --- |
| TPS02_Arabidopsis thaliana | Y (0.973 \| 1-26) | - | - |
| TPS02_Cr1504 | Y (0.992 \| 1-47) | - | - |
| TPS02_Cg926 | Y (0.993 \| 1-34) | - | - |
| TPS03_Arabidopsis thaliana | - | Y (0.894 \| 1-45) | Y (RRMVIHALEMPYHRRVGR) |
| TPS03_Cr1504 | - | - | Y (RKMVIHALEMPYHRRVGR,RRMIGVAWDDLNLKRICR) |
| TPS03_Cg926 | - | - | Y (RKMVIHALEMPYHRRVGR,RRMIGVAWDDLNLKRICR) |

**Table S7.**
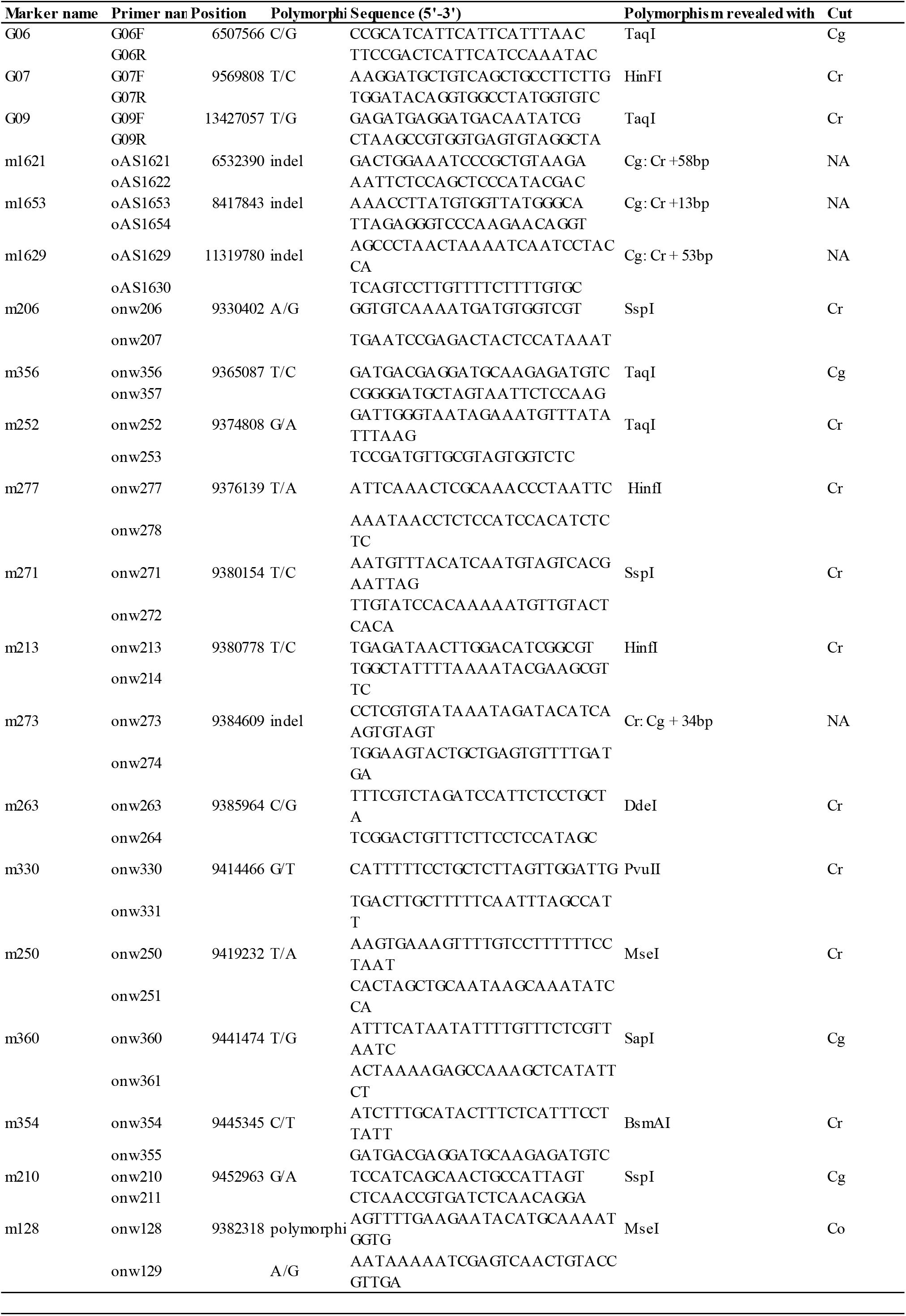
Primer sequences and information about PCR-based markers used for mapping

**Table S8.**
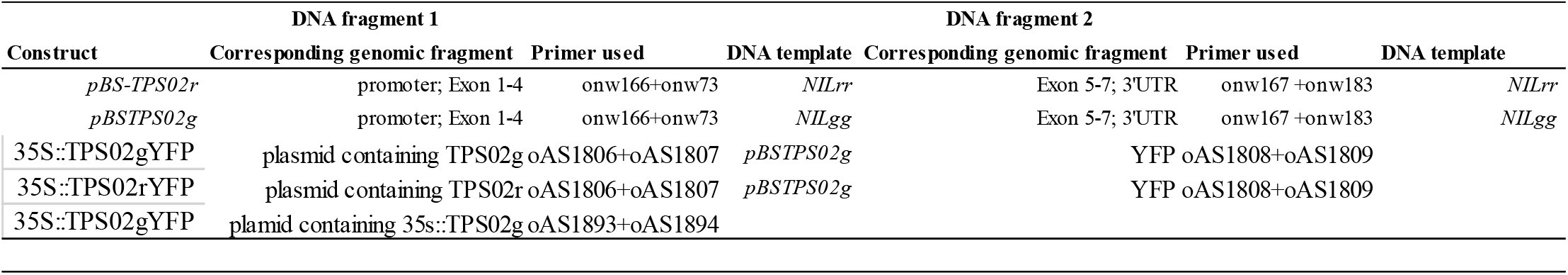
PCR fragments used in the molecular cloning reaction.

**Table S9.**
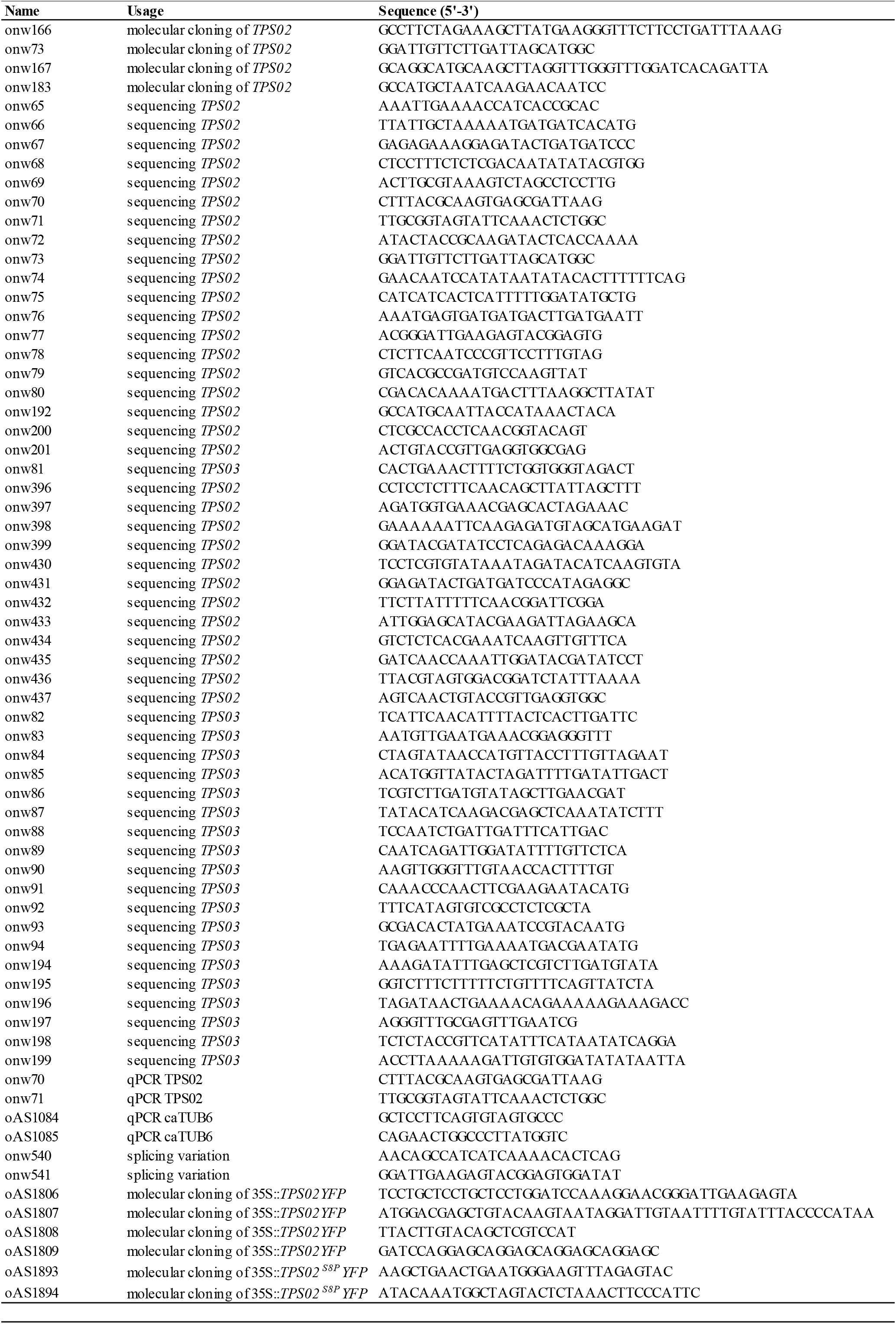
Primer sequences used in this study.

## Notes

### Competing Interest Statement

The authors have declared no competing interest.

